# Transcriptional plasticity of melanin-concentrating hormone neurons in the medial preoptic area of lactating mouse dams

**DOI:** 10.64898/2026.05.12.724557

**Authors:** Allan dos Anjos-Monteiro, Jozélia Gomes Pacheco Ferreira, Victor Hugo dos Santos-Affonso, Elizabeth Chamiec-Case, Laura E. Mickelsen, Behrokh Marzban Abbasabadi, Carol Fuzeti Elias, Alexander C. Jackson, Jackson Cioni Bittencourt

## Abstract

The medial preoptic area (MPOA) is a central hub for maternal behavior, integrating hormonal and sensory signals to coordinate adaptive postpartum responses. Although melanin-concentrating hormone (MCH) neurons are well characterized in the lateral hypothalamus, their identity and functional engagement within MPOA circuits remain poorly defined. Here, through integrative reanalysis of a publicly available single-cell RNA sequencing dataset of the mouse MPOA (GSE295610), we identify two transcriptionally distinct *Pmch*-expressing neuronal populations. Both populations are GABAergic and emerge prominently during mid to late lactation. Lactation is characterized by significant upregulation of *Pmch* and coordinated enrichment of neuropeptidergic and hormone-responsive genes, including islet amyloid polypeptide (*Iapp*), prodynorphin (*Pdyn*), and prolactin receptor (*Prlr*). Independent single-cell gene expression profiling of FACS-isolated GAD67-GFP neurons from the MPOA further corroborated these findings, confirming the selective emergence of *Pmch* expression during lactation and its co-expression with neuropeptidergic and hormone-responsive genes. NeuroEstimator-based activity inference demonstrates increased predicted neuronal activity in lactating females, while pseudotime reconstruction reveals a lactation-associated transcriptional shift toward later trajectory states. hdWGCNA analysis identified gene co-expression modules significantly enriched during lactation. Regulatory network inference using SCENIC further revealed activation of activity-dependent transcriptional regulons, including cyclic AMP-responsive element-binding protein 3-like 1 (*Creb3l1*), early growth response 1 (*Egr1*), and FBJ osteosarcoma oncogene (*Fos*). These transcriptional programs converge on gene networks associated with synaptic plasticity, regulation of neurogenesis, and broader mechanisms of neuronal plasticity. Notably, these *Pmch* populations were not annotated in the original study, underscoring the power of systems-level reanalysis to uncover previously unrecognized components of maternal circuitry. Together, our findings provide single-cell evidence that MCH-expressing neurons in the MPOA undergo state-dependent transcriptional reorganization during lactation, suggesting a dynamic role for MCH signaling in postpartum neuroendocrine plasticity.

---

Maternal behavior is an evolutionarily conserved set of coordinated responses arising from the interaction between a mother and her offspring. These responses ensure offspring survival by providing nutrition, warmth, protection, and social stimulation during early development (1–4). In mammals, these behaviors are broadly divided into two functional domains: maternal care and maternal aggression (5,6). In rodents, maternal care includes stereotyped repertoires such as licking and grooming, nursing, pup retrieval, and nest building, whereas maternal aggression consists of defensive responses toward potential threats, including attacks and threat displays directed at intruders (7–10)

The neural control of maternal behavior relies on an interconnected network that integrates endocrine signals with sensory information originating from the offspring. At the center of this network, the medial preoptic area of the hypothalamus (MPOA) acts as a key hub co-ordinating maternal responses (11–15). Seminal studies demonstrate that MPOA lesions or reversible neuronal inactivation impairs maternal behavior during early post-partum period (16–19).

The functional contribution of the MPOA appears to be state-dependent, as manipulation of this region during later stages of lactation can produce distinct behavioral outcomes (20). This suggests that MPOA circuitry undergoes dynamic reorganization across the maternal period. MPOA neurons also integrate somatosensory and olfactory cues from the offspring, allowing mothers to co-ordinate physiological and behavioral adaptations required for their care (21–23)

Consistent with this integrative role, single-cell transcriptomic studies have revealed that the MPOA contains a highly heterogeneous cellular landscape composed of multiple transcriptionally distinct neuronal populations (24–28). Although these approaches have greatly expanded our understanding of MPOA cellular diversity, the transcriptional identity and state-dependent regulation of specific neuronal populations engaged during lactation remain incompletely understood.

Among hypothalamic neuromodulatory systems potentially involved in maternal adaptations, melanin-concentrating hormone (MCH) neurons have emerged as important regulators of motivated and social behaviors (29,30). MCH is derived from the pro-melanin-concentrating hormone (*Pmch*) gene, which encodes the precursor peptide that gives rise to MCH, primarily in the lateral hypothalamic area (LHA) (31–33). Notably, transient expression of MCH has also been reported in the MPOA during lactation in females (34,35), suggesting that this neuropeptidergic system may participate in maternal state–dependent adaptations within MPOA circuits regulating approach–avoidant behaviors during the postpartum period (36,37).

Given our limited understanding of the neurochemical and functional organization of MCH-expressing neurons in the MPOA (MPOA^*Pmch*^) and the pronounced cellular heterogeneity of this region, defining the MCH-related transcriptional programs at single-cell resolution will offer new insights into the neuronal plasticity and the state-dependent remodeling of MPOA circuitry. Here, we performed an integrative reanalysis of a previously published scRNA-seq dataset from the MPOA of virgin and lactating female mice (*GSE295610*). By combining unsupervised clustering, differential expression analysis (DEGs), and network-based approaches, we identified transcriptionally distinct neuronal populations and characterized the transcriptional landscape of MPOA^*Pmch*^ neurons associated with maternal adaptations.

## Results

### Transcriptomic characterization of the MPOA dataset reveals a heterogeneous and hormonally responsive cellular landscape

To delineate the cellular architecture of the MPOA, we reanalyzed a previously published sc-RNA-seq (*GSE295610*) (24), comprising 10x Genomics profiles from the MPOA of adult female C57BL/6 mice (14 weeks old). The dataset includes samples from sated virgin females and lactating mothers collected at postpartum days 13–15, with three pooled animals per condition. For the present analysis, only MPOA-derived cells from these two groups were included. (**Fig. 1A**). Quality-control metrics, including the number of detected genes, total UMI counts, mitochondrial transcript percentage, and ribosomal transcript percentage, were examined to ensure data quality (**Fig. S1A–D**).

**Fig. 1.**
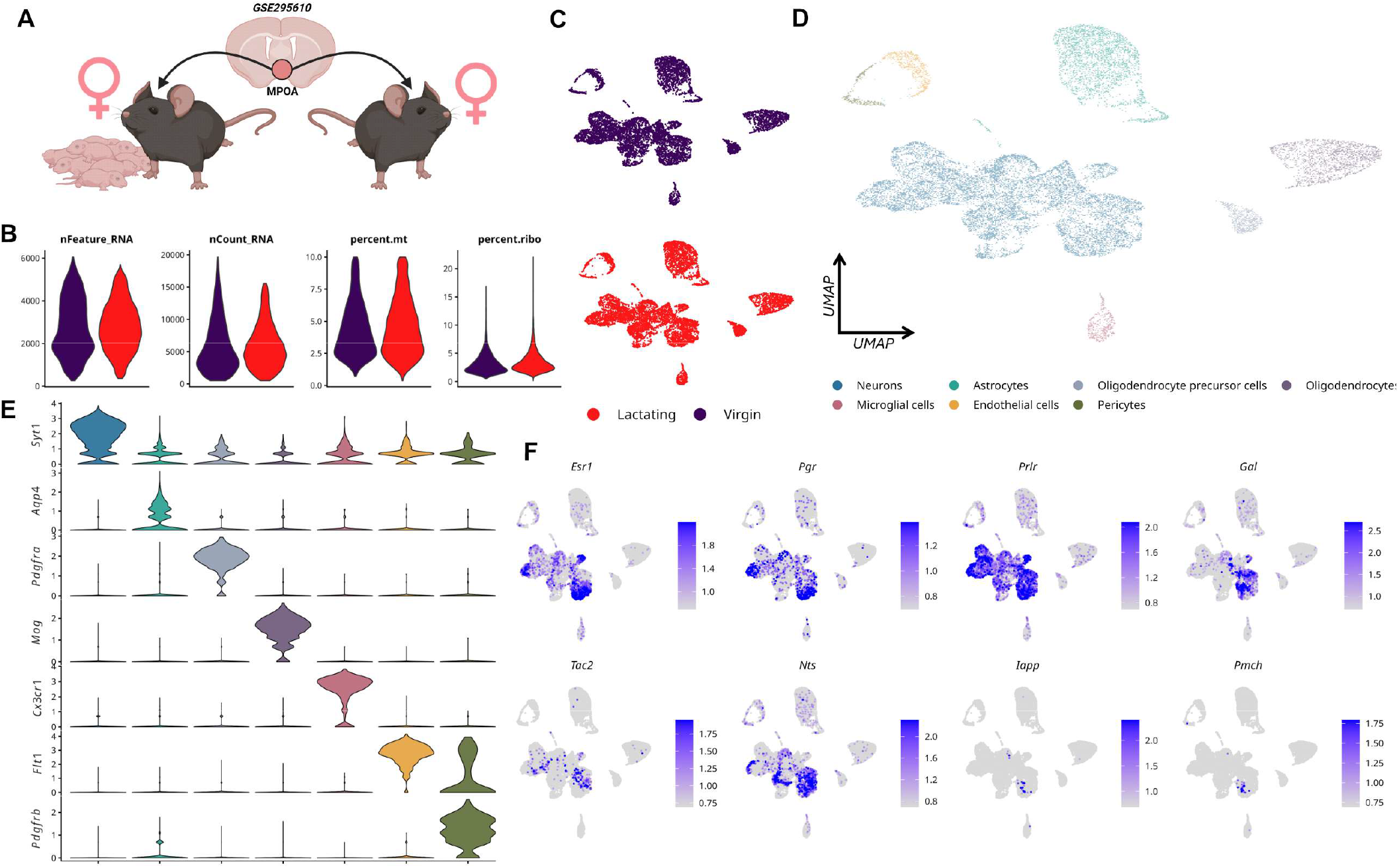
Single-cell transcriptomic landscape of the MPOA across maternal states reveals cellular heterogeneity and neuroendocrine specialization. **A**. Schematic representation of the experimental design showing integration of single-cell RNA-seq data from virgin and lactating females to characterize the cellular architecture of the MPOA (GSE295610). **B**. Quality control metrics across conditions, including number of detected genes (*nFeature_RNA*), total counts (*nCount_RNA*), and percentage of mitochondrial and ribosomal transcripts, indicating comparable data quality between virgin and lactating groups. **C**. UMAP projection colored by physiological state reveals overlapping global cellular distributions between virgin and lactating animals, indicating that maternal state does not broadly alter overall cellular composition of the MPOA. **D**. Cell-type annotation identifies major populations within the MPOA, including neurons, astrocytes, oligodendrocyte precursor cells, oligodendrocytes, microglia, endothelial cells, and pericytes, highlighting the cellular diversity of this region. **E**. Violin plot visualization of canonical marker genes confirms cell-type identity, including neuronal (*Syt1*), astrocytic (*Aqp4*), oligodendrocyte lineage (*Pdgfra, Mog*), microglial (*Cx3cr1*), endothelial (*Flt1*), and pericyte (*Pdgfrb*) markers, supporting robust annotation of MPOA cell populations. **F**. Feature plots of key neuroendocrine and MPOA-associated genes reveal spatially restricted expression patterns across neuronal populations, including hormone receptors (*Esr1, Pgr, Prlr*), neuropeptides (*Gal, Tac2, Nts, Pmch*), and metabolically relevant signaling molecules (*Iapp*). These patterns highlight the presence of hormonally responsive and neuropeptidergic neuronal populations within the MPOA, consistent with its role as an integrative hub coordinating endocrine signals and maternal behavioral outputs.

Relationships between sequencing depth and gene detection, as well as mitochondrial and ribosomal transcript content, were further evaluated to define appropriate filtering thresholds (**Fig. S1E–F**). Predicted doublets were subsequently identified and removed before downstream analyses **(Fig. S1G**). Following these filtering steps, cells from both conditions exhibited comparable transcriptomic complexity and mitochondrial transcript content, indicating consistent data quality and the absence of major technical biases between groups (**Fig. 1B)**.

UMAP visualization revealed extensive overlap between cells derived from virgin and lactating animals within a shared transcriptional space, supporting effective integration and minimizing potential condition-specific batch effects (**Fig. 1C**). The proportional representation of the identified cell populations was comparable between conditions, indicating similar cellular composition across datasets (**Fig. S1H–I**).

Unsupervised clustering analysis revealed a heterogeneous cellular landscape comprising neural and non-neural populations. Major cell classes included neurons, astrocytes, oligodendrocyte precursor cells (OPCs), mature oligodendrocytes, microglial cells, endothelial cells, and pericytes (**Fig. 1D; Table S1**). Cell-type identities were assigned based on the expression of established canonical marker genes, following annotation frameworks previously defined in single-cell studies of the hypothalamus and preoptic area (38,39) (see **Table S1** for the complete list of markers used for annotation).

Cell-type annotations were validated through the expression of canonical marker genes. Neuronal populations displayed robust expression of synaptotagmin-1 (*Syt1*), astrocytes expressed aquaporin-4 (*Aqp4*), OPCs expressed platelet-derived growth factor receptor alpha (*Pdgfra*), oligodendrocytes expressed myelin oligodendrocyte glycoprotein (*Mog*), microglial cells expressed C-X3-C motif chemokine receptor 1 (*Cx3cr1*), endothelial cells expressed Fms-related receptor tyrosine kinase 1 (*Flt1*), and pericytes expressed platelet-derived growth factor receptor beta (*Pdgfrb*) **(Fig. 1E)**. Together, these canonical markers confirmed the accurate classification of the principal cellular populations within the dataset.

Importantly, feature expression visualization of classical MPOA-associated and neuroendocrine-related genes demonstrated the presence of hormonally responsive neuronal populations. Key regulators of reproductive and parental circuits, including estrogen receptor alpha (*Esr1*), progesterone receptor (*Pgr*), prolactin receptor (*Prlr*), galanin (*Gal*), and neurokinin B (*Tac2*), were enriched within defined neuronal subpopulations, consistent with the well-established role of the MPOA as a central hub controlling reproductive physiology and maternal behavior (**Fig. 1F**). In addition, neuropeptidergic genes such as neurotensin (*Nts*), islet amyloid polypeptide (*Iapp*), and *Pmch* highlighted further transcriptional specialization among neuronal subsets, suggesting the presence of distinct neuroendocrine programs within MPOA circuitry.

### Emergence and functional activation of a Lama2/Esr1 neuronal subpopulation during lactation

To further resolve neuronal diversity within the MPOA, we performed a focused reclustering of the neuronal population. UMAP reduction revealed multiple transcriptionally distinct neuronal subclusters organized according to top marker gene expression profiles (**Fig. 2A; Fig. S2A;** see the gene markers for each subcluster in **Table S2**), highlighting molecular signatures associated with distinct hypothalamic neuronal identities (**Fig. 2B**). To further contextualize these neuronal populations, label transfer analysis using the Moffitt high-resolution MER-FISH atlas was performed (27), enabling cross-dataset correspondence between the identified MPOA neuronal subclusters and previously annotated hypothalamic cell populations **(Fig. S2B)**.

**Fig. 2.**
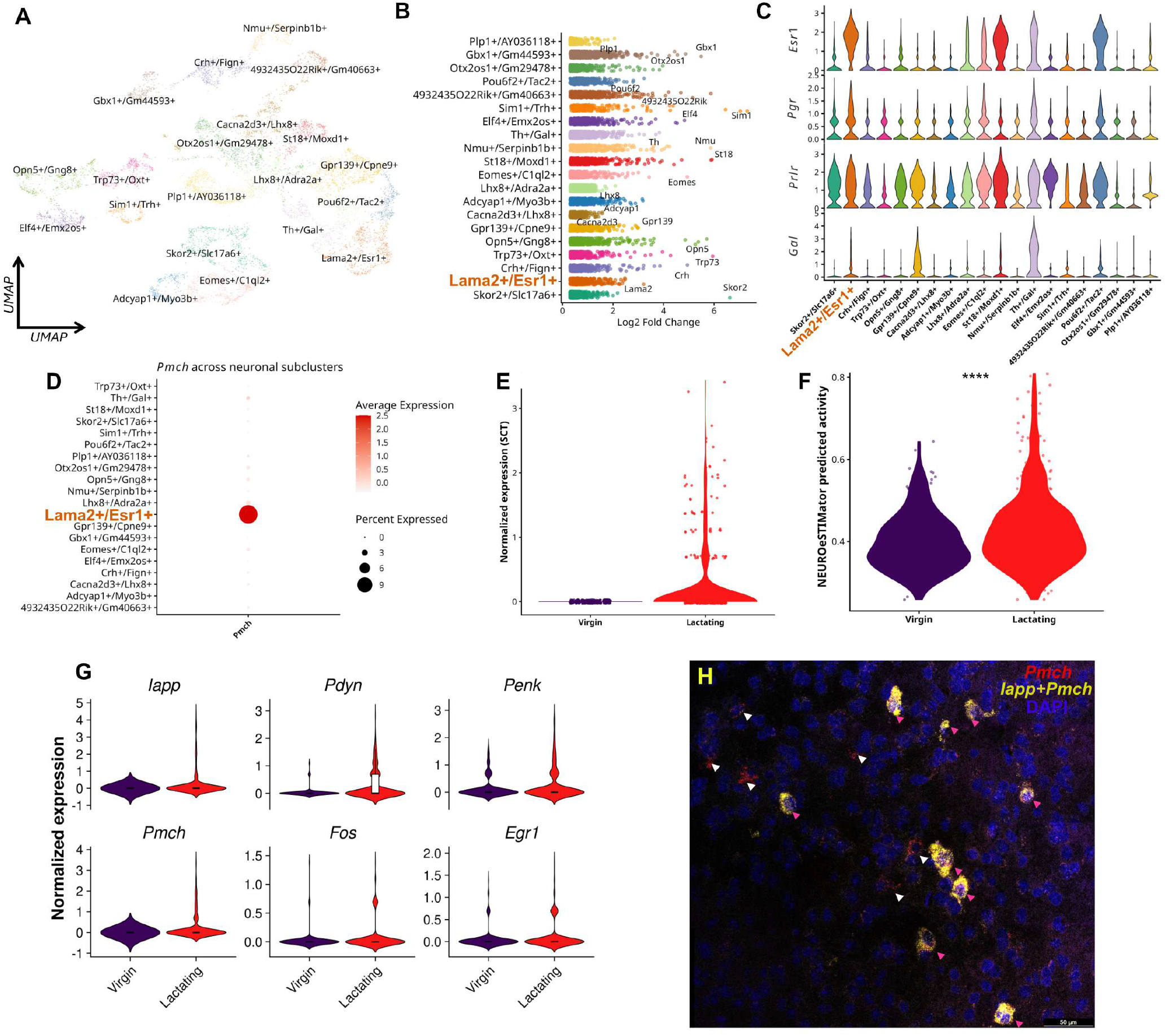
Transcriptional identification and functional characterization of *Pmch* neuronal populations in the MPOA. **A**. UMAP projection of neuronal subclusters within the MPOA reveals transcriptionally distinct populations, including a discrete MPOA^*Lama2*/*Esr1*^ cluster enriched for *Pmch* expression, suggesting selective localization of MCH-related transcriptional programs within defined neuronal subtypes. **B**. Differential gene expression analysis across neuronal clusters highlights marker genes defining each population, supporting the molecular segregation of MPOA neuronal identities. **C**. Violin plots showing expression of key neuroendocrine markers across neuronal subclusters, including hormone receptors (*Esr1, Pgr, Prlr*) and neuropeptides (*Gal*), indicating the presence of hormonally responsive neuronal populations within the MPOA. **D**. Dot plot visualization of *Pmch* expression across neuronal subclusters demonstrates selective enrichment within specific populations, supporting a restricted distribution of MCH neurons within MPOA circuitry. **E**. Expression levels of *Pmch* across physiological states reveal increased expression in lactating animals, indicating maternal state–dependent transcriptional regulation. **F**. NEUROeSTIMator analysis revealed significantly increased predicted neuronal activity in lactating animals, indicating enhanced functional engagement of MPOA neuronal populations during the maternal state.**G**. Violin plots showing expression patterns of representative genes associated with lactation-related *Pmch* neuronal populations, including *Iapp, Pdyn, Penk, Pmch, Fos*, and *Egr1*. These analyses indicate enrichment of neuropeptidergic and activity-associated transcriptional programs during lactation. **H**. RNAscope validation confirming the presence of *Pmch/Iapp* co-expressing neurons in the MPOA of lactating mice. Pink arrowheads indicate cells showing *Pmch/Iapp* co-localization (yellow signal). Nuclei were counterstained with DAPI (blue). Scale bar: 50 μm.

These markers included transcription factors, ion channels, neuropeptides, and signaling molecules, reflecting the functional heterogeneity of MPOA neurons. Notably, several genes previously associated with hypothalamic and MPOA circuits were enriched across neuronal clusters, including the transcription factors single-minded homolog 1 (*Sim1*), POU class 6 homeobox 2 (*Pou6f2*), and T-box brain transcription factor 2 (*Eomes*), the neuropeptide corticotropin-releasing hormone (*Crh*), and the catecholamine synthesis enzyme tyrosine hydroxylase (*Th*). Additional markers such as LIM homeobox 8 (*Lhx8*) and neuromedin U (*Nmu*) further highlighted the molecular diversity of neuronal populations within the MPOA (**Fig. 2B**).

We next examined the distribution of hormone-responsive genes across neuronal subclusters and the neurotransmitter identity of MPOA neurons. Visualization of gene expression revealed heterogeneous expression profiles for key regulators of reproductive and parental circuits, including *Esr1, Pgr, Prlr*, and *Gal* (**Fig. 2C**).

To further characterize the synaptic identity of these neuronal populations, we analyzed the expression of canonical markers of excitatory and inhibitory neurotransmission. Feature visualization revealed that a subset of MPOA neurons expressed *Slc17a6* (MPOA^Vglut2^), indicative of a glutamatergic phenotype, whereas the majority of neuronal populations expressed *Slc32a1* (MPOA^Vgat^), consistent with a predominantly GABAergic profile (**Fig. S2C-D**). Together, these findings demonstrate that MPOA neuronal subclusters exhibit heterogeneous hormone receptor expression while encompassing both excitatory and inhibitory neuronal populations, highlighting the complex molecular and synaptic organization of this hypothalamic region.

To investigate the distribution of MPOA^*Pmch*^, we mapped its expression across neuronal clusters. This analysis revealed a selective enrichment within the MPOA^*Lama2/Esr1*^ neuronal cluster (**Fig. 2D, Fig. S2E**), identifying a transcriptionally defined subpopulation of MPOA^*Pmch*^ neurons. Quantitative comparison between physiological conditions showed that *Pmch* expression was markedly increased in lactating females relative to virgin animals among MPOA^*Lama2/Esr1*^ neurons (**Fig. 2E**).

To determine whether this transcriptional shift was associated with functional activation of neuronal populations, we performed activity inference using NeuroeS-TIMator within the MPOA^*Lama2/Esr1*^ neuronal cluster, a deep learning–based framework that integrates activity-dependent transcriptional signatures to estimate neuronal activation from single-cell transcriptomic data (40). This analysis revealed significantly higher predicted neuronal activity scores in lactating animals compared with virgin females (**Fig. 2F**), indicating increased functional activation of this neuronal population during lactating maternal state.

We next investigated the transcriptional context surrounding *Pmch* expression within the MPOA *Lama2/Esr1* neuronal cluster to identify genes associated with this putative neuropeptidergic population. Expression analyses revealed coordinated enrichment of neuropep-tide-related genes, including *Iapp*, prodynorphin (*Pdyn*), and proenkephalin (*Penk*), together with activity-associated genes such as FBJ osteosarcoma oncogene (*Fos*) and early growth response 1 (*Egr1*) **(Fig. 2G)**, suggesting that these neurons belong to a broader neuropeptidergic and activity-responsive transcriptional program within the MPOA.

To validate the transcriptional findings at the spatial level, we performed RNAscope *in situ* hybridization to examine the distribution of *Pmch* and *Iapp* transcripts within the MPOA. Consistent with the single-cell transcriptomic analysis, RNAscope revealed discrete neuronal populations expressing *Pmch, Iapp*, as well as neurons co-expressing both transcripts within the MPOA (**Fig.2H**). Notably, all *Iapp* neurons co-expressed *Pmch*, whereas a subset of *Pmch* neurons did not express *Iapp*, indicating that *Iapp*-expressing cells represent a transcriptionally defined subpopulation within the broader *Pmch* neuronal pool.

These findings support the transcriptional heterogeneity observed in the single-cell dataset and provide spatial validation for the predicted neuropeptidergic identities of *Pmch*-associated neuronal populations.

### Distinct *Pmch*-expressing neuronal subtypes defined by *Tmem45a*/*Ror1* and *Gm33782*/*4933425B07Rik* molecular identities

To further investigate the molecular heterogeneity of MPOA^*Lama2/Esr1*^, we performed a focused analysis of MPOA enriched for *Pmch* expression. Reclustering of neuronal populations revealed several transcriptionally distinct subtypes organized according to characteristic gene expression profiles (**Fig. 3A**). Visualization of cells by physiological condition demonstrated that both virgin and lactating animals contributed to these neuronal populations, although their distribution across the transcriptional landscape differed, suggesting state-dependent recruitment of specific neuronal identities (**Fig. 3B**).

**Fig. 3.**
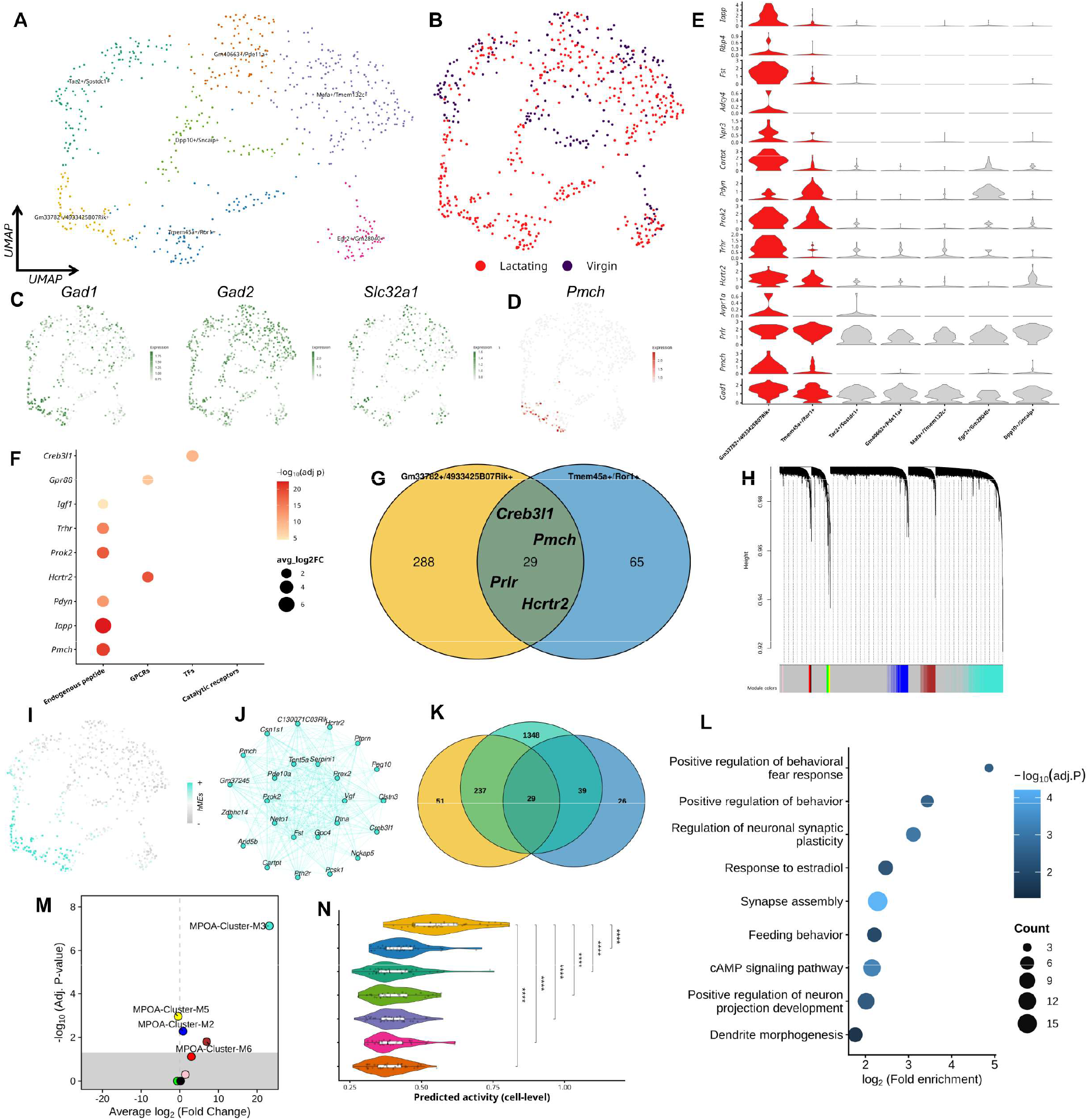
Molecular characterization of *Pmch* neuronal subpopulations in the MPOA^*Lama2/Esr1*^. **A**. UMAP embedding of MPOA neuronal subclusters annotated according to top marker genes. Two transcriptionally distinct neuronal populations are defined by *Tmem45a*/*Ror1* and *Gm33782*/*4933425B07Rik* identities. **B**. UMAP colored by physiological condition (Virgin vs Lactating), illustrating lactation-associated redistribution within neuronal territories. **C**. Feature plots of inhibitory markers (*Gad1, Gad2, Slc32a1*) demonstrate that both *Pmch*-expressing clusters belong to a GABAergic lineage. **D**. Feature plot of *Pmch* expression reveals localization within two spatially and transcriptionally distinct neuronal domains corresponding to *Tmem45a*/*Ror1* and *Gm33782*/*4933425B07Rik* clusters. **E**. Violin plots showing neuropeptide receptors and hormone-responsive genes across neuronal subclusters, indicating differential regulatory landscapes between the two *Pmch* populations. **F**. Dot plot summarizing enrichment of endocrine receptors, GPCRs, and neuropeptide-related genes, highlighting subtype-specific signaling repertoires. **G**. Venn diagram showing overlap between *Tmem45a*/*Ror1* and *Gm33782*/*4933425B07Rik* cluster markers. Shared genes include *Creb3l1, Pmch, Prlr*, and *Hcrtr2*, suggesting a partially convergent neuroendocrine program despite distinct molecular backbones. **H**. hdWGCNA module eigengenes distribution analysis identifying co-expression modules enriched in *Pmch* clusters. **I**. UMAP projection of module activity scores demonstrates spatial restriction of lactation-associated transcriptional programs. **J**. Regulatory network centered on *Pmch*, showing predicted transcription factor–target interactions and co-regulated genes. **K**. Multi-set intersection analysis integrating cluster markers and the M-3 module. **L**. Gene Ontology enrichment of the shared core reveals overrepresentation of behavioral regulation, synaptic plasticity, response to estradiol, feeding behavior, and cAMP signaling. **M**. Volcano plot of module-level DMEs highlighting lactation-associated transcriptional remodeling. **N**. Predicted neuronal activity scores across clusters show increased activity in *Pmch*-expressing subtypes during lactation. * *p* < 0.05, ** *p* < 0.01, *** *p* < 0.001.

To define the neurotransmitter identity of these neuronal populations, we examined the expression of canonical inhibitory markers. Feature visualization revealed strong expression of the GABAergic markers glutamate decarboxylase 1 (*Gad1*), glutamate decarboxylase 2 (*Gad2*), and the vesicular GABA transporter *Slc32a1* across neuronal clusters (**Fig. 3C**), indicating that these MPOA^*Lama2*/*Esr1*^ populations primarily belong to inhibitory neuronal circuits within the MPOA.

Mapping of MPOA^*Pmch*^ expression across the neuronal landscape revealed that *Pmch* transcripts were restricted to a discrete subset of neurons (**Fig. 3D**). Detailed inspection of marker gene expression profiles identified two transcriptionally distinct *Pmch*-expressing neuronal populations, defined by two top marker combinations (gene markers are provided in **Table S3**): *Tmem45a/Ror1* and *Gm33782/4933425B07Rik* molecular signatures (**Fig. 3E, Table S6**). To further resolve the transcriptional organization of these populations, *Pmch*-expressing neurons were subsetted and independently re-clustered (**Fig. S3A**)

Notably, these neurons exhibited enrichment of several genes for neuropeptides and receptors, including *Iapp, Pdyn*, cocaine- and amphetamine-regulated transcript prepropeptide (*Cartpt*), prokineticin 2 (*Prok2)*, thyrotropin-releasing hormone receptor (*Trhr*), hypocretin receptor 2 (*Hcrtr2*), and arginine vasopressin receptor 1A (*Avpr1a*), together with the core marker *Pmch* (**Fig. 3E**). These transcriptional signatures remained distinct following focused reanalysis of the *Pmch* neuronal subset (**Fig. S3B–C**). Scatterplot analyses further supported the segregation of these neuronal populations based on differential marker associations with *Pmch* expression (**Fig. S3D**). These findings reveal previously unrecognized neurochemical features of MPOA *Pmch* neurons, suggesting functional specialization within this neuronal population. This GABAergic identity was further supported by fluorescence *in situ* hybridization analyses (**Fig. S4A–B**).

Gene distribution analysis based on International Union of Basic and Clinical Pharmacology (IUPHAR) classification revealed enrichment of multiple neuropeptide and signaling-related genes within these MPOA^*Pmch*^ neuronal populations, including genes linked to neuroendocrine signaling and neuronal communication (**Fig. 3F**). Enriched categories included endogenous peptides, G protein–coupled receptors (GPCRs), transcription factors, and catalytic receptors (**Table S4**).

Intersection analysis of subtype-specific markers revealed a shared transcriptional core comprising genes such as cyclic AMP-responsive element-binding protein 3-like 1 (*Creb3l1*), *Pmch, Prlr*, and *Hcrtr2*, indicating a conserved neuroendocrine regulatory module present in both neuronal subtypes (**Fig. 3G, Table S5**).

To investigate the co-expression networks associated with *Pmch*-expressing neurons, we performed weighted gene co-expression network analysis using hdWGCNA. An appropriate soft-thresholding power was selected to construct the gene co-expression network (**Fig. S4C**). The resulting hierarchical clustering dendrogram revealed the organization of genes into discrete co-expression modules across MPOA neuronal populations (**Fig. 3H**). Module connectivity was further assessed by calculating module eigengene-based connectivity (kME) values, which identified the most highly connected genes within each module (**Fig. S4D**).

Gene co-expression network analysis revealed tightly connected transcriptional modules centered on *Pmch* and associated neuropeptidergic genes, particularly within Module-M3 (**Fig. 3I-J**) (all identified modules and their associated genes are provided in **Table S6**). The distribution of additional modules and their corresponding network structures further illustrates the transcriptional heterogeneity across MPOA neuronal subtypes (**Fig. S4F-G**). Intersection analysis of network modules revealed a subset of genes shared between the *Tmem45a/Ror1* and *Gm33782/4933425B07Rik clusters*, suggesting partially overlapping regulatory programs despite their distinct molecular signatures (**Fig. 3K**) (list of intersecting genes is provided in **Table S7**).

Functional enrichment analysis revealed significant overrepresentation of biological processes related to regulation of neuronal synaptic plasticity (GO:0048168), synapse assembly (GO:0007416), positive regulation of neuron projection development (GO:0010976), and dendrite morphogenesis (GO:0048813) (**Fig. 3L**). Additional enrichment was observed for signaling and hormonal response pathways, including the cAMP signaling pathway (mmu04024) and response to estradiol (GO:0032355). Behavior-related processes were also significantly enriched, including positive regulation of behavior (GO:0048520), positive regulation of behavioral fear response (GO:2000987), and feeding behavior (GO:0007631) (**Table S8** provides the complete list of enriched pathways).

Finally, differential module eigengene expression (*DME*) analysis across MPOA^*Lama2/Esr1*^ neuronal clusters demonstrated that *Pmch*-expressing populations correspond to distinct transcriptional states, characterized by the upregulation of Module-M3 genes associated with neuropeptide signaling and hormonal responsiveness (**Fig. 3M**).

Consistent with these transcriptional signatures, activity inference analysis further revealed significantly increased predicted neuronal activity within specific MPOA^*Lama2/Esr1*^ neuronal clusters enriched for *Pmch* expression (**Fig. 3N**) (statistical comparisons supporting these analyses are provided in **Table S9**).

Together, these results indicate that *Pmch*-expressing neurons within the MPOA^*Lama2/Esr1*^ population are not homogeneous, but instead comprise two transcriptionally distinct GABAergic neuronal subtypes, defined by *Tmem45a/Ror1* and *Gm33782/4933425B07Rik* molecular identities. Despite their distinct molecular signatures, these populations share a conserved neuroendocrine transcriptional core, while maintaining subtype-specific regulatory programs.

### *Pmch* neuronal populations occupy a central position in MPOA^*Lama2/Esr1*^ communication networks

To determine how *Pmch*-expressing neuronal populations are integrated within MPOA circuitry, we performed cell cell communication analysis restricted to the MPOA^*Lama2/Esr1*^ neuronal population using CellChat.

Global network visualization revealed a highly interconnected communication landscape among neuronal subtypes, with *Pmch*-enriched populations (*Tmem45a/Ror1* and *Gm33782/4933425B07Rik*) exhibiting extensive incoming and outgoing interactions in a lactation state-dependent manner (**Fig. 4A–B; Fig. S5A**). These populations occupy a highly connected position within the network, consistent with a central role in local signal integration (see **Table S10** for a complete list of cellular interactions).

**Fig. 4.**
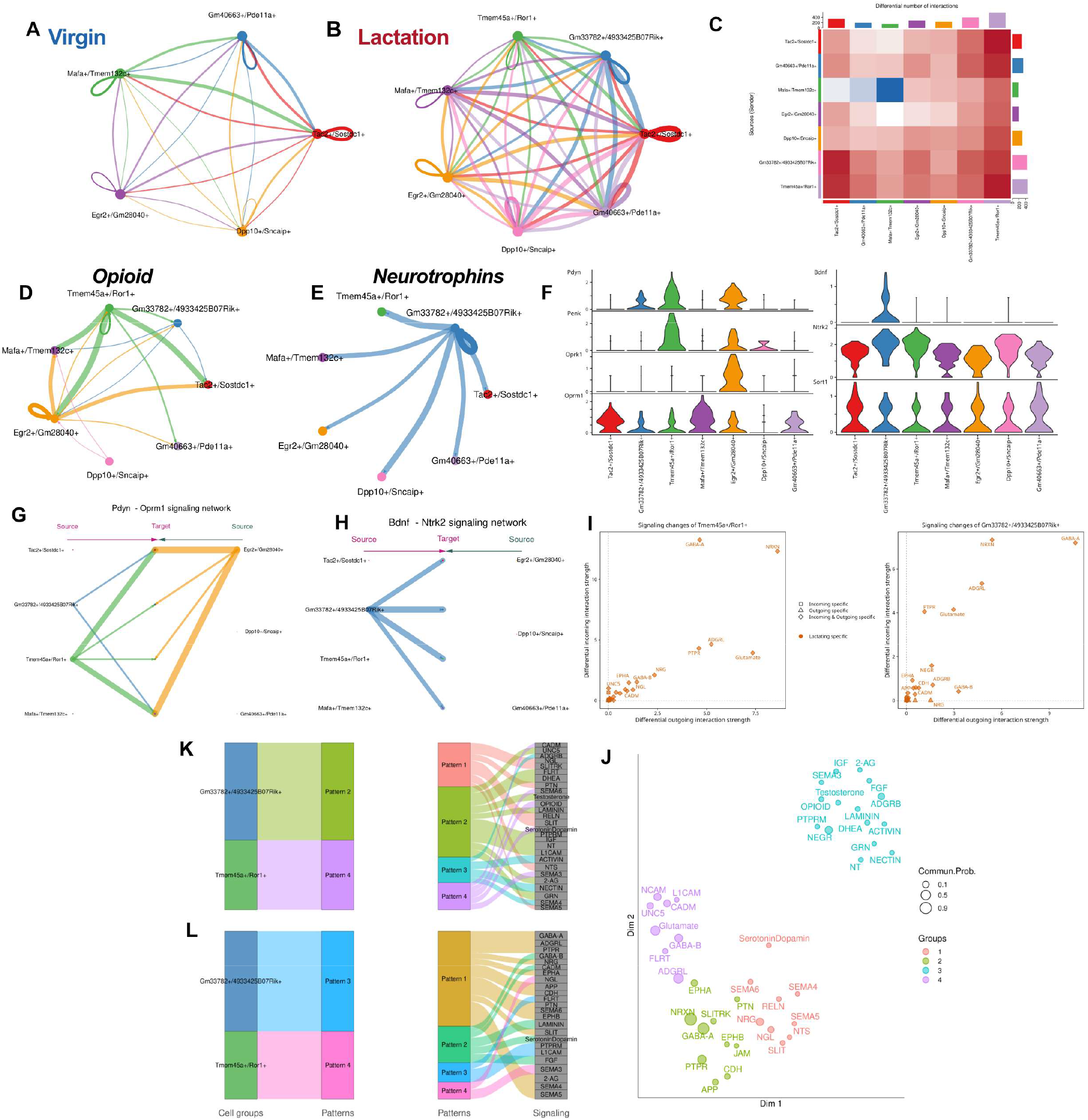
Cell–cell communication analysis reveals state-dependent integration of *Pmch* neurons within MPOA networks. **A–B**. Global communication networks inferred by CellChat showing interactions among *Lama2*/*Esr1* neuronal subtypes under virgin and lactation conditions, high-lighting extensive incoming and outgoing signaling associated with *Pmch*-enriched populations (*Tmem45a/Ror1* and *Gm33782/4933425B07Rik*). **C**. Heatmap of relative differential interaction strength across neuronal populations, illustrating the contribution of each subtype to overall information flow. **D–E**. Representative signaling pathways, including opioid (*Pdyn–Oprm1*) and neurotrophin (*Bdnf–Ntrk2*) networks, showing structured and directed interactions centered on *Pmch* subpopulations. **F**. Expression patterns of key ligand–receptor components across neuronal subtypes, supporting inferred communication networks. **G–H**. Source–target interaction networks for selected pathways, highlighting directionality of signaling between neuronal populations. I**–J**. Differential interaction analysis showing changes in outgoing and incoming signaling strength, with pathways clustering into functional groups and a dominant module enriched for lactation-associated signaling within *IGF, RELN, NT/NTS*, semaphorin (*SEMA4/5*), *FGF*, and laminin pathways. **K–L**. Alluvial plots illustrating the mapping between neuronal subpopulations, signaling patterns, and pathway identities, revealing organized communication programs and their association with distinct functional signaling modules.

Quantitative analysis of interaction strength showed that *Pmch* subpopulations contribute substantially to overall information flow within the MPOA^*Lama2/Esr1*^ network (**Fig. 4C; Fig. S5B**), indicating that these neurons are integral components of the communication architecture.

Pathway-level analysis identified engagement of neuropeptidergic and growth-related signaling systems. Opioid (*Pdyn–Oprm1*) and neurotrophin (*Bdnf–Ntrk2*) signaling form structured interaction networks centered on *Pmch* subpopulations, together with prominent GABA-A and GABA-B signaling (**Fig. 4D–E, G–I; Fig. S5C–D**), consistent with coordinated modulatory and inhibitory functions.

Expression analysis confirmed that ligand–receptor components underlying these interactions are differentially enriched across *Pmch* subtypes (**Fig. 4F**), supporting the inferred communication structure. *NEGR*-associated signaling, along with GABA-A and *NRXN*-mediated interactions, further support roles in synaptic organization,inhibitory transmission, and network connectivity within the MPOA.

Differential interaction analysis showed that signaling pathways cluster into functional groups. At the systems level, signaling pathways organize into distinct communication programs, with modules enriched for adhesion-related signaling (*NRXN, NCAM, CADM*), neurotransmission (GABA-A/B, glutamate), and neuropeptidergic and growth-related pathways (*RELN, IGF, NTS*), which occupy segregated regions of the signaling space and align with lactation-associated profiles (**Fig. 4K–L**).

A dominant module enriched during lactation included *IGF, RELN, NT/NTS*, semaphorin (*SEMA4/5*), *FGF*, and laminin signaling (**Fig. 4J; Fig. S5E–F**), indicating coordinated recruitment of pathways associated with circuit remodeling. Pattern selection analysis supported the robustness of these communication programs (**Fig. S5G–H**). These pathways also showed increased outgoing signaling from *Pmch* subpopulations during lactation, consistent with a more active signaling role. Together, these results position *Pmch* neurons as central nodes within MPOA communication networks, with enhanced signaling engagement during lactation.

### Single-cell profiling of GAD67-GFP neurons reveals a lactation-dependent neuropeptidergic population in the MPOA

As an orthogonal approach to experimentally define the neurochemical identity of MPOA *Pmch* neurons, we performed single-cell profiling using reverse transcription quantitative polymerase chain reaction (RT–qPCR). MPOA tissue was isolated using anatomically guided micriopunches from GAD67-GFP mice **(Fig. 5A–C′)**. Adult animals (~3 months old) of both sexes and reproductive conditions were used, including males, virgin females, and lactating females, with lactating females collected at postpartum day 19 (PPD19).

**Fig. 5.**
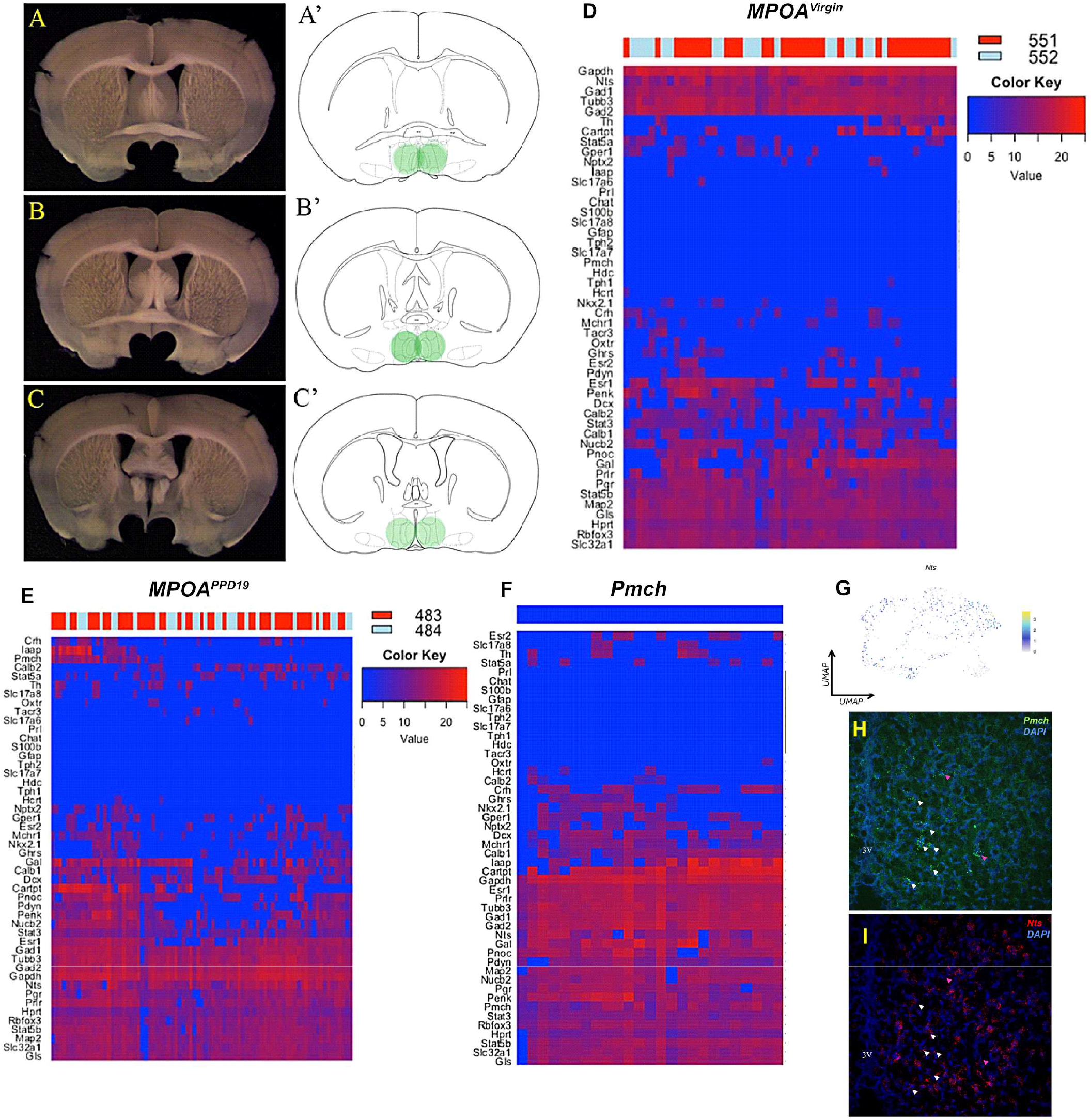
Neurochemical characterization and spatial validation of *Pmch*-expressing neurons in the MPOA using GAD67-GFP mice. **A–C**. Representative coronal brain sections showing the anatomical localization of the MPOA across rostrocaudal levels. **A′–C′**. Schematic representations indicating the region of interest (green) used for micropunch isolation. **D**. Heatmap showing gene expression profiles of MPOA neurons isolated from GAD67-GFP mice at postpartum day 19 (PPD19), highlighting neurochemical heterogeneity and enrichment of neuropeptidergic and GABAergic markers. **E-F**. Heatmap of *Pmch* neurons identified from FACS-purified GAD67-GFP cells, displaying a distinct transcriptional profile enriched for inhibitory and neuropeptidergic genes. **G-I**. RNAscope showing *Pmch* (green) and *Nts* (red) expression in the MPOA, with DAPI (blue). Arrowheads indicate cells co-expressing Pmch and Nts. The third ventricle (3V) and anterior commissure (ac) is indicated for anatomical reference.

Following tissue dissociation, single EGFP neurons were isolated by fluorescence-activated cell sorting (FACS) (**Fig. S6A–D**). Cells were sorted at single-cell resolution and initially screened by RT–qPCR for *Gapdh, Nts*, and *Pmch* expression (**Fig. S6E-F)**

Subsequently, FACS-purified single neurons were subjected to high-throughput gene expression profiling using *TaqMan* assays in a 48.48 dynamic array on a *Biomark HD* system (Fluidigm), enabling the analysis of 48 selected genes across neurochemically defined populations (**Fig. S6G;** see **Table S11**). This strategy enabled the selective identification and molecular characterization of neurochemically defined neuronal populations.

Consistent with this approach, *Pmch* expression was not detected in virgin females (**Fig. 5D**), whereas it was predominantly observed in lactating animals (**Fig. 5E**). Quantitative screening further indicated that *Pmch* expression defines a small but enriched neuronal subset during lactation, supporting a state-dependent emergence of this population.

Single-cell qPCR profiling revealed that *Pmch* neurons exhibit a complex neurochemical signature. In addition to GABAergic markers, these cells coexpress multiple neuropeptides, including *Iapp, Crh, Nptx2, Gal, Nts, Cartpt, Pdyn, Penk*, and *Nucb2*, as well as hormone-responsive genes such as *Esr1, Pgr, Prlr*, and members of the *Stat* family (**Fig. 5F**). Conversely, these neurons show little to no expression of canonical glutamatergic, monoaminergic, or glial markers, reinforcing their identity as neuropeptidergic GABAergic neurons.

Consistent with these findings, fluorescence *in situ* hybridization confirmed the spatial localization of *Pmch*-expressing neurons within the MPOA and demonstrated a high degree of co-expression with neurotensin, similar to that observed in MPOA^Lama2/Esr1^ clusters (**Fig. 5G-I**), supporting the integration of these neurons within a broader neuropeptidergic network.

### Lactation-associated transcriptional dynamics and regulatory programs suggest neuroplastic remodeling within MPOA *Pmch* neurons

To investigate transcriptional dynamics associated with maternal state, we performed pseudotime trajectory analysis of MPOA^*Lama2/Esr1*^ neuronal subpopulations. Reconstruction of the transcriptional trajectory revealed a continuous progression of cellular states across the MPOA^*Lama2/Esr1*^ manifold (**Fig. 6A**). Cells from lactating animals were preferentially distributed along later pseudotime positions, whereas neurons from virgin animals were enriched in earlier states, indicating a lactation-associated shift in transcriptional programs. Quantitative comparison confirmed a significant displacement toward later pseudotime stages in lactating animals (**Fig. 6B**), supporting a state-dependent transcriptional transition within MPOA^*Lama2/Esr1*^ neurons.

**Fig. 6.**
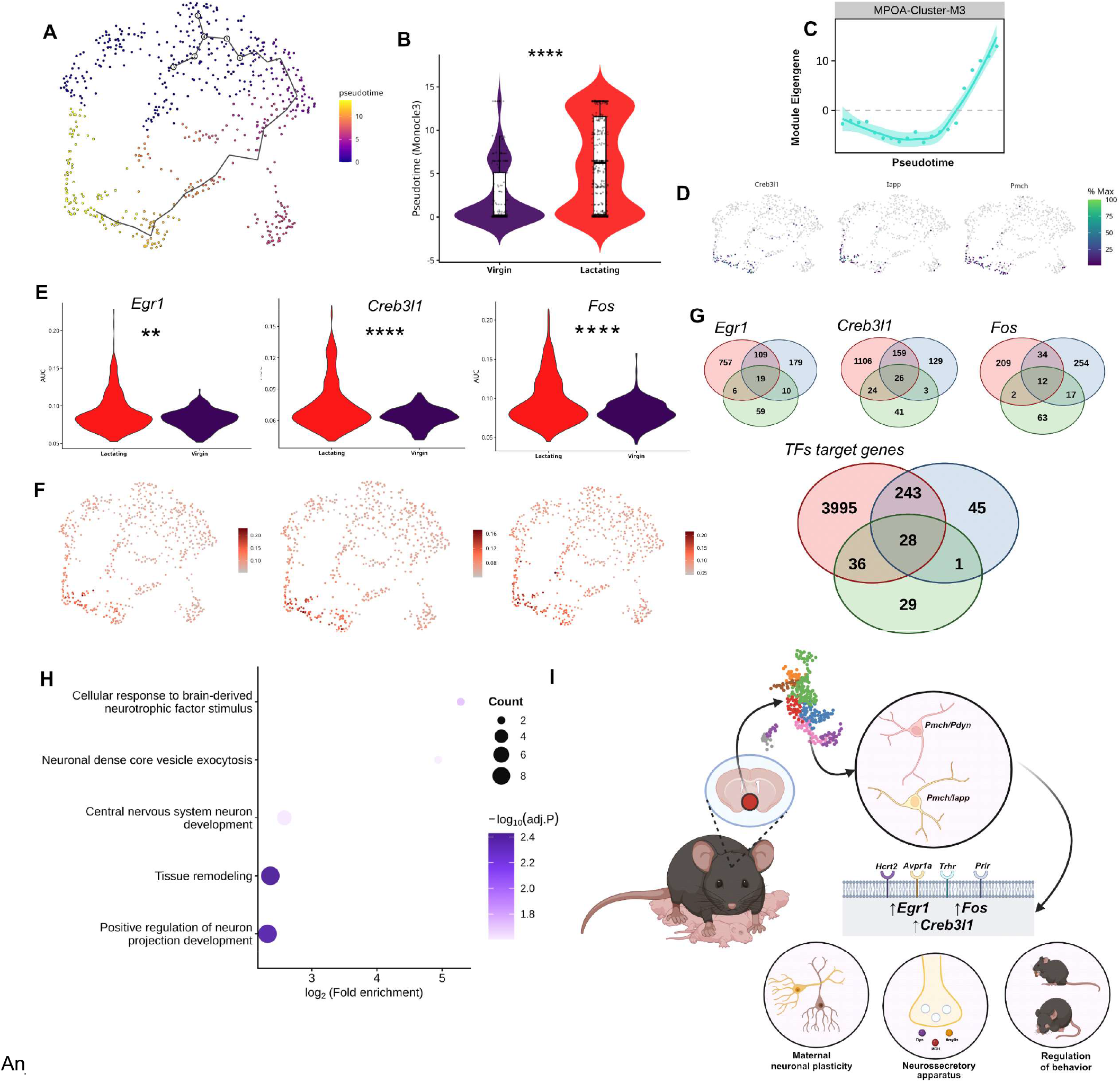
Transcriptional dynamics and regulatory programs associated with *Pmch* neuronal activation during lactation. **A**. Pseudotime trajectory analysis of MPOA neurons reveals a continuous transcriptional progression across cellular states, suggesting dynamic remodeling of neuronal identity. **B**. Distribution of pseudotime values across physiological conditions shows a significant shift toward advanced transcriptional states in lactating animals, indicating state-dependent progression rather than transient activation. **C**. Module eigengene analysis of the MPOA-associated module (M3) demonstrates increased activity along pseudotime, consistent with coordinated transcriptional activation during lactation. **D**. Feature plots of representative genes (*Creb3l1, Iapp, Pmch*) across the trajectory illustrate progressive upregulation along pseudotime, supporting their involvement in state-dependent transcriptional programs. **E**. Violin plots showing increased expression of activity-dependent transcription factors (*Egr1, Creb3l1, Fos*) in lactating animals, indicating enhanced transcriptional activation. **F**. UMAP visualization of transcription factor activity scores further supports increased regulatory activity in lactating conditions. **G**. Intersection analysis of transcription factor target genes reveals both shared and unique regulatory programs for *Egr1, Creb3l1*, and *Fos*, identifying a partially convergent transcriptional network in *Pmch* neuronal clusters. **H**. Functional enrichment analysis of transcription factor targets highlights pathways related to neuronal plasticity, including BDNF response, dense-core vesicle exocytosis, neuronal differentiation, and tissue remodeling, consistent with experience-dependent circuit reorganization. **I**. Schematic model summarizing the proposed role of *Pmch* neurons in the MPOA. Hormonal and sensory inputs converge onto *Pmch*-expressing neuronal populations, engaging transcriptional regulators (*Creb3l1, Egr1, Fos*) and downstream signaling pathways to drive neuronal plasticity, neurosecretory function, and regulation of maternal behavior. * *p* < 0.05, ** *p* < 0.01, *** *p* < 0.001

To characterize gene expression changes along this trajectory, we examined the spatial distribution of key neuropeptidergic and regulatory genes. Feature visualization revealed dynamic expression patterns of *Creb3l1, Iapp*, and *Pmch* across pseudotime, closely mirroring the M3 module eigengene dynamics (**Fig. 6C–D; Fig. S7A**), indicating coordinated regulation of genes associated with neuropeptide signaling, hormonal responsiveness, and neuronal plasticity (see **Table S12**).

Given the involvement of activity-dependent transcriptional programs, we next assessed transcription factor activity using the SCENIC framework. Regulon activity analysis revealed increased activity of *Egr1, Creb3l1*, and *Fos* in lactating animals compared with virgins (**Fig. 6E**), consistent with enhanced transcriptional activation of *Pmch*-associated neuronal populations. Additional transcription factors predicted to bind regulatory regions of *Pmch* were also identified (**Fig. S7B–E**), suggesting potential upstream regulatory inputs. Spatial mapping of transcription factor activity further demonstrated enrichment across specific regions of the neuronal manifold (**Fig. 6F; Fig. S7F-I**), indicating structured regulatory engagement.

To investigate regulatory convergence, we examined overlap among transcription factor target genes. Intersection analysis revealed a shared set of downstream targets for *Egr1, Creb3l1*, and *Fos* (**Fig. 6G**), supporting coordinated transcriptional regulation within *Pmch*-expressing neurons. Broader integration of regulon targets further identified a core set of shared genes across transcription factor networks (**Fig. S7J–M; Table S13**), indicating convergence onto common regulatory modules.

Functional enrichment analysis of genes associated with these regulatory programs revealed significant overrepresentation of biological processes related to neuronal development and plasticity, including positive regulation of neuron projection development (GO:0010976), central nervous system neuron development (GO:0021954), tissue remodeling (GO:0048771), cellular response to brain-derived neurotrophic factor stimulus (GO:1990416), and neuronal dense core vesicle exocytosis (GO:0099011) (**Fig. 6H**). These pathways are consistent with structural and functional adaptations occurring within MPOA circuits during lactation (**a complete list of enriched pathways is provided in Table S14**).

Together, these findings indicate that lactation is associated with coordinated transcriptional progression, regulatory network convergence, and activation of gene programs linked to neuronal plasticity and neuropeptide signaling within *Pmch*-associated MPOA neurons. These dynamics support a model in which maternal state drives structured transcriptional remodeling of MPOA circuits, integrating hormonal and activity-dependent signals to promote adaptive behavioral outputs (**Fig. 6I**).

## Discussion

The MPOA integrates endocrine, sensory, and physiological signals to orchestrate maternal behavior (3,4,41) Although this region is a well-established hub of maternal circuitry, the molecular diversity of neuronal populations underlying postpartum adaptations remains incompletely defined.

Through integrative reanalysis of scRNA-seq data (24). We identified two transcriptionally distinct *Pmch* neuronal populations within the MPOA that emerge during mid-to late lactation. These findings expand the current framework of MPOA cellular heterogeneity and position MCH signaling as a previously underappreciated component of transcriptional and circuit-level remodeling in maternal networks.

At the cellular level, the MPOA is predominantly composed of GABAergic neurons and exhibits strong sensitivity to reproductive hormones, including estrogen, progesterone, and prolactin (42–46). In addition, the MPOA contains multiple neuropeptidergic and receptor-defined neuronal populations implicated in parental behavior, including *Gal, Nts, Calcr*, and *Oxtr* (47–51), reinforcing its role as a neurochemical integration hub linking environmental inputs to behavioral outputs.

MCH neurons have been extensively characterized in the LHA, where they regulate energy balance, arousal, sleep, and motivated behaviors (52–55). In contrast, their role within the MPOA has remained poorly defined and largely restricted to late lactation (35). In mice, MCH neurons are detectable in the MPOA during lactation, with immunoreactivity emerging around PPD19 (56). Here, enrichment of *Pmch* expression within MPOA^*Lama2*/*Esr1*^ subclusters suggests that MCH signaling is embedded within *Esr1*-dependent maternal circuits, extending prior anatomical observations into a transcriptional framework.

Within MPOA^*Lama2/Esr1*^ neurons, *Iapp* (amylin) emerged as a key maternally regulated gene. Previous studies identified *Iapp* as one of the most strongly induced transcripts in the MPOA of lactating rats (57), and largely segregated from MCH populations in lactating dams (58). MCH expression in the MPOA increases during lactation and continues to rise through the late postpartum period, when maternal responsiveness naturally declines. Consistent with this, local MCH administration in the MPOA selectively impairs active components of maternal behavior, such as pup retrieval and nest building, without affecting nursing (59). Together, these observations raise the possibility that MCH contributes to avoidance-related behavioral responses emerging during the late postpartum period in rats.

In contrast, our scRNA-seq analysis identified MPOA neurons co-expressing *Pmch* and *Iapp*, and RNAscope validation confirmed substantial overlap between these markers in lactating mice. Thus, *Iapp* expression is robustly induced around parturition and remains elevated during lactation, with amylin-expressing MPOA neurons strongly activated by pup exposure and closely associated with maternal behavioral expression (60). Moreover, amylin signaling regulates affiliative contact-seeking behavior in female mice (47). Together, these findings suggest that *Pmch* and *Iapp* may contribute to complementary aspects of maternal adaptation, integrating maternal engagement with its progressive modulation across the postpartum period.

Additional co-expressed neuropeptides further position these neurons within established MPOA maternal circuits. Galanin (*Gal*) marks a core MPOA neuronal population required for parental behavior and coordinates distinct motor, motivational, and social components through projection-specific pathways (61,62). Neurotensin (*Nts*) signaling is also upregulated in the MPOA during the postpartum period and has been implicated in the modulation of maternal responsiveness (63)

Consistent with this framework, independent single-cell gene expression profiling of GABAergic MPOA neurons further corroborated the selective emergence of *Pmch* expression during lactation and its coordinated co-expression with hormone-responsive genes.

*Pmch* expression in the MPOA was resolved into two molecularly distinct neuronal populations characterized by *Tmem45a/Ror1* and *Gm33782/4933425B07Rik* signatures. Both populations exhibited a GABAergic profile, consistent with evidence that MPOA MCH neurons co-express *Slc32a1* (29), and expressed *Prlr*, in line with prolactin-responsive neuronal populations engaged during lactation (64).

Despite this transcriptional divergence, intersection analysis revealed a shared transcriptional core comprising *Pmch, Pdyn, Prlr*, and *Hcrtr2*, defining a conserved neuroendocrine regulatory module. The inclusion of *Hcrtr2* is consistent with evidence that hypocretin signaling within the MPOA promotes maternal behavior (65,66), linking this module to the regulation of maternal responses.

Moreover, *Pdyn* and *Cartpt* have been described in male mice MPOA neuronal populations and are implicated in the regulation of motivational and affective behaviors (67) suggesting that this shared transcriptional program may integrate neuropeptidergic systems involved in the modulation of both social and internal states. Co-expression network analysis revealed enrichment of pathways associated with synaptic plasticity and neuronal structural remodeling, including synapse assembly and dendrite morphogenesis, indicating experience-dependent reorganization. Concomitant enrichment of hormonal and intracellular signaling pathways further suggests integration of endocrine inputs with transcriptional and synaptic regulation (68).

In parallel, enrichment of behavior-related processes indicates that these neurons may coordinate motivational, affective, and metabolic components of maternal responses. Together, these findings support a model in which MPOA *Pmch* neurons integrate hormonal, motivational, and emotional signals to shape adaptive maternal behavioral states, consistent with the well-established plasticity of the maternal brain (69,70).

Experimental evidence further supports dynamic regulation of MCH neurons in the MPOA. Hormonal manipulations indicate sensitivity to ovarian steroids, while correlations with litter size and pup-derived cues highlight integration of endocrine and sensory inputs in lactating rat dams (71,72).

Multiple analytical approaches indicate functional engagement of these neurons during lactation. NeuroeS-TIMator analysis revealed increased predicted activity in lactating females, while pseudotime analysis demonstrated progression toward later transcriptional states, consistent with sustained state transitions rather than transient activation. Together, these findings suggest that *Pmch* neurons are dynamically recruited and stabilized within maternal circuits, consistent with extensive evidence demonstrating activity-dependent plasticity within the MPOA during maternal behavior, as reflected by *Fos*-like expression patterns driven by pup-related sensory inputs and maternal experience (73,74).

Regulatory network analysis further supports this interpretation. SCENIC identified increased activity of TFs including *Creb3l1, Egr1*, and *Fos. Creb3l1* is linked to *Esr1*-dependent transcriptional programs in females (26), whereas *Fos* and *Egr1* represent activity-dependent immediate early genes induced by pup exposure, indicating layered temporal regulation of transcriptional responses (75).

Beyond their role as activity markers, *Egr1* has been identified as a regulator of neuronal chromatin accessibility and structural plasticity, driving experience-dependent transcriptional remodeling in a sex-dependent manner (76). This suggests that activity-dependent gene induction in the MPOA may not only reflect transient neuronal activation but also contribute to longer-term adaptations in circuit function during lactation.

Consistent with this framework, previous studies have shown that neurotensin-dependent *Egr1* activity is differentially regulated across reproductive states, with reduced sensitivity observed during the postpartum period, further supporting dynamic reconfiguration of activity-dependent transcriptional responses in maternal circuits (77,78).

At the circuit level, cell–cell communication analysis revealed that *Pmch* neuronal populations are embedded within structured MPOA signaling networks and occupy highly connected positions within the *Lama2/Esr1* microenvironment. These neurons engage neuropeptidergic, inhibitory, and adhesion-related pathways, including *Pdyn–Oprm1*, neurotrophin (*Bdnf–Ntrk2*), GABA-A/B, consistent with roles in synaptic organization and circuit integration (79–83). Notably, lactation is associated with enrichment of extracellular signaling pathways, such as IGF, RELN, NTS, and semaphorin families, accompanied by increased outgoing signaling from *Pmch* subpopulations, indicating a shift toward a more active role in MPOA network dynamic.

In accordance with this line of data, enrichment of pathways related to neuronal development, CNS differentiation, BDNF signaling, and dense-core vesicle exocytosis indicates coordinated structural and functional remodeling of MPOA circuits, likely contributing to stabilization of maternal behavioral states (84).This coordinated remodeling is likely to enhance synaptic plasticity and peptidergic signaling within MPOA networks, thereby reinforcing the persistence and adaptability of maternal behavioral responses across the postpartum period.

## Conclusion

In summary, we identify previously uncharacterized *Pmch* neuronal populations in the MPOA that undergo coordinated transcriptional activation during mid to late lactation. These findings support a model in which MCH neurotransmission contributes to state-dependent remodeling of maternal circuits and reveals a previously unrecognized role for *Pmch* neurons in postpartum neuroendocrine plasticity. More broadly, this work underscores the power of targeted, systems-level reanalysis to uncover previously unrecognized neuronal identities. Collectively, we establish a comprehensive, multi-level characterization of MPOA *Pmch* neurons, integrating transcriptional, regulatory, and functional dimensions to define their role within maternal neuroendocrine circuits.

## Acknowledgements

This work was supported by the Brazilian Coordenação de Aperfeiçoamento de Pessoal de Nível Superior (CAPES). We are grateful for the support provided by Fundação de Amparo à Pesquisa do Estado de São Paulo (FAPESP). J.C.B. is an Investigator with the Conselho Nacional de Desenvolvimento Científico e Tecnológico (CNPq; grants #404806/2021-0 and #302203/2022-2). We thank members of the Laboratory of Chemical Neuroanatomy at the University of São Paulo for valuable discussions and technical support. We are grateful to Carol Fuzeti Elias, Alexander C. Jackson, Jackson Cioni Bittencourt. This study was supported by FAPESP (grants #2023/02531-5, #2025/13454-7, and #2026/04203-3, 2018/04659-0) and CAPES.

## Author contributions

A.d.A.M. conceived the study, data generation, performed data analysis and interpretation, and wrote the manuscript. J.G.P.F. contributed to experimental procedures and data acquisition. V.H.S.A. contributed to computational analyses and data interpretation. E.C.-C. and L.E.M. contributed to experimental work and data generation. B.M. contributed to data analysis and interpretation. C.F.E., A.C.J., and J.C.B. provided conceptual guidance, contributed to data interpretation, and revised the manuscript. All authors approved the final version of the manuscript.

## Materials and Methods

### Animals

Experiments were performed using genetically modified mouse lines to characterize the neurochemical identity of *Pmch*-expressing neurons in the MPOA. GAD67–GFP mice were used to identify and isolate GABAergic neuronal populations. For neurochemical and single-cell analyses, adult mice (~3 months old) of both sexes and across reproductive states were included, comprising males, virgin females, and lactating females. Lactating animals were analyzed at postpartum day 19 (PPD19), a stage associated with peak maternal neuro-endocrine adaptations. Animals were maintained under standard laboratory conditions with ad libitum access to mice chow and water.

### Micropunch isolation and FACS sorting of GAD67-GFP neurons

To isolate GABAergic neurons from the MPOA, female GAD67-GFP mice were euthanized at PPD19 by decapitation. Brains were rapidly removed and sectioned coronally (225 μm thickness) using a vibratome. The ventro-medial medial preoptic area (vmMPOA) was identified based on anatomical landmarks, i.e., the third ventricle and the crossing of the anterior commissure, and isolated using a 1-mm diameter micropunch. Microdissected tissue was enzymatically and mechanically dissociated, triturated, and filtered to obtain a single-cell suspension. Cells were subjected to fluorescence-activated cell sorting (FACS) using a BD FACSAria II system. GFP^+^ neurons were identified based on fluorescence and forward/side scatter properties, and singlets were gated to ensure single-cell resolution. Individual cells were sorted into 96-well plates containing lysis buffer, immediately frozen on dry ice, and stored at −80 °C until further processing.

### Single-cell RT-qPCR and microfluidic gene expression analysis

Sorted single cells were subjected to reverse transcription followed by cDNA pre-amplification. Gene expression profiling was performed using TaqMan Gene Expression Assays in a 48.48 dynamic array format on the Biomark HD microfluidic system (Fluidigm). A curated panel of 48 genes was selected to capture neurotransmitter identity, neuropeptidergic signaling, and hormone responsiveness (**Table S15**), encompassing markers of GABAergic transmission, neuropeptides, receptors, and intracellular signaling pathways. Initial quality control and cell classification were performed by RT–qPCR targeting *Gapdh, Pmch*, and *Nts*, enabling identification of viable cells and neurochemical subtypes. Subsequent analyses were restricted to cells meeting these criteria.

### Fluorescence *in situ* hybridization (RNAscope)

To validate gene co-expression at the spatial level, fluorescence *in situ* hybridization (FISH) was performed using the RNAscope platform (ACD). Coronal brain sections containing the MPOA were processed with probes targeting *Pmch, Slc32a1, Nts*, and *Iapp*. Tissue sections were prepared following manufacturer guidelines, including fixation, permeabilization, and probe hybridization steps optimized for single-molecule detection of mRNA transcripts. Signal amplification was achieved through a series of sequential hybridization steps, enabling high-sensitivity detection with minimal background. Fluorescent signals were visualized using confocal microscopy, allowing high-resolution imaging of individual cells and precise assessment of transcript co-localization. Z-stack imaging and optical sectioning were used to confirm cellular co-expression within the same neuronal profiles. Co-localization analyses were performed to determine the spatial overlap between neuropeptidergic and GABAergic markers, providing anatomical validation of the molecular signatures identified through transcriptomic and single-cell approaches.

### Analysis of transcriptional data from BioMark HD

The cycle threshold values (Ct) for each marker were obtained through specialized software, inverted and then used in the construction of a log-based scale for analysis of gene expression. Only samples with demonstrated control gene synthesis (*Gapdh* and *Hprt*) were used in the analysis. For correlation analyses, the Ward method was used for hierarchical grouping, followed by principal component analysis (PCA).

### Dataset Acquisition

Single-cell RNA sequencing data were obtained from the Gene Expression Omnibus (GEO) under accession number *GSE295610*. The dataset comprises 10x Genomics Chromium profiles generated from MPOA and arcuate nucleus (ARC) of 14-week-old female C57BL/6 mice across four experimental conditions: sated virgins (VS), fasted virgins (VF), sated lactating mothers (MS), and fasted lactating mothers (MF), with three pooled animals per group(24). Lactating samples were collected at postpartum days 13–15. For the present study, only MPOA-derived samples from VS and MS groups were included. VF and MF samples were excluded to avoid confounding effects of acute metabolic stress and to isolate transcriptional adaptations specifically associated with the lactational state under baseline feeding conditions. Raw count matrices were downloaded and reprocessed *de novo* in R using Seurat to ensure analytical consistency across all preprocessing and downstream analyses.

### Quality Control and Filtering

Quality control (QC) was performed using Seurat version 5 (85) in R (4.5.2). Per-cell metrics included the number of detected genes (nFeature_RNA), total UMI counts (nCount_RNA), percentage of mitochondrial transcripts (percent.mt), and percentage of ribosomal transcripts (percent.ribo). Mitochondrial content was calculated and ribosomal content was quantified from canonical ribosomal protein genes. To determine the mitochondrial filtering threshold, retention curves were generated across candidate cutoffs (3–25%) and the inflection point was used to guide selection; cells with percent.mt > 10% were excluded. To identify high-UMI and high-ribosomal outliers while accounting for potential condition-specific distributional differences, robust thresholds were computed independently within VS and MS groups using a median absolute deviation (MAD)-based approach.

Upper bounds for nCount_RNA and percent.ribo were defined as the median plus five times the MAD. Cells were retained if they satisfied all of the following criteria: 200 ≤ nFeature_RNA ≤ 10,000, nCount_RNA below the condition-specific MAD threshold, percent.mt ≤ 10%, and percent.ribo below the condition-specific MAD threshold. This strategy minimized the inclusion of low-complexity cells, stressed cells, and extreme high-count outliers while preserving biologically meaningful state-dependent variation.

### Doublet detection, integration, clustering, and cell-type annotation

After initial QC filtering, putative doublets were identified and removed using scDblFinder (86). The filtered Seurat object was converted to a SingleCellExperiment and scDblFinder was run with sample stratification by condition (virgins and lactating), returning per-cell doublet classifications and scores. Cells classified as singlets were retained for all subsequent analyses. Singlet cells were normalized and variance-stabilized using SCTransform, with percent.mt included as a covariate. PCA was performed on the SCTransform assay, and the top PCs (n = 25) were used as input for Harmony to correct for condition-associated technical variation while preserving biological structure. Nearest-neighbor graphs were constructed using Harmony embeddings, and unsupervised clustering was performed across a range of resolutions to evaluate cluster stability; a single resolution was selected for downstream analyses and reporting were generated from Harmony-corrected dimensions for visualization and presentation.

Cell-type annotation was performed at the major neuronal cell types using ScType (87). ScType marker gene sets were provided via a curated brain reference spreadsheet, and ScType scores were computed from the scaled expression matrix to assign a best-matching sub-type label to each cluster. For each cluster, the top-scoring ScType label was selected as the subtype annotation, and low-confidence assignments were conservatively labeled as “Unknown” based on an empirical score-to-cell-number criterion. These subtype annotations were then used for downstream marker discovery and visualization.

### Neuronal subset extraction, reprocessing, and sub-clustering

Following generation of the Harmony-integrated MPOA object annotations, neuronal cells were extracted. The neuronal subset was then reprocessed independently to maximize resolution within neuronal transcriptional space. Specifically, the RNA assay was set as default and the data were variance-stabilized using SCTransform (v2 flavor), while regressing out mitochondrial transcript burden (percent.mt). Dimensionality reduction was performed by principal component analysis (PCA) on the SCTransform assay (50 PCs computed), and the number of PCs used for graph-based analyses was selected based on inspection of the elbow plot (25 PCs used for downstream steps). A k-nearest neighbor graph was constructed using the selected PCs, followed by unsupervised clustering across a grid of resolutions (0.1–1.0) to assess granularity and cluster stability. Dimensional embeddings (UMAP) were generated from the PCA space for visualization, and cluster structure as well as sample composition were visualized using UMAP colored by clustering assignment and by original sample identity.

To further evaluate clustering robustness and separation across resolutions, silhouette analysis was performed using PCA embeddings derived from the SCTransform assay. Average silhouette widths were computed for each clustering resolution based on Euclidean distances in reduced-dimensional space, allowing quantitative assessment of intra-cluster cohesion and inter-cluster separation. These metrics were used as an additional diagnostic measure to support cluster resolution selection and evaluate the stability of transcriptionally defined neuronal populations.

### Differential expression, marker discovery, and cluster labeling within neurons

To identify robust marker genes defining neuronal subclusters, differential expression analyses were performed in a one-versus-rest framework for each cluster at the selected clustering resolutions (0.5 for the global neuronal subset and 0.8 for focused subcluster analyses). Marker testing was performed using the MAST algorithm (88), considering only positively enriched genes and applying minimum expression frequency and effect-size thresholds (min.pct = 0.10–0.25; log fold-change threshold = 0.25–0.30). Statistical significance was determined following multiple-testing correction (adjusted p value < 0.05).

### High-Dimensional Weighted Gene Co-Expression Network Analysis (hdWGCNA)

Gene co-expression networks were constructed within the neuronal subset using hdWGCNA (89). Genes were selected based on fractional expression (≥ 5% of cells), and metacells were generated within each cluster and sample using k-nearest neighbor aggregation (k = 5; minimum 15 cells per group) to reduce sparsity and improve network stability. The normalized metacell expression matrix derived from the SCT assay was used for network construction. Signed weighted gene co-expression networks were generated using Pearson correlation and topological overlap (TOM). Soft-thresholding (β = 8) power was selected according to scale-free topology criteria.

Modules were detected using dynamic tree cutting (deepSplit = 4; minimum module size = 50 genes) and merged at a cut height of 0.25. Module eigengenes (MEs) were computed and projected to the full single-cell dataset, with harmonization across samples to control for batch structure. Intramodular connectivity (kME) was calculated to quantify gene-module membership strength. Differential module eigengene (DMEs) analysis between biological groups was performed using Mann-Whitney non-parametric testing. Module structure, eigengene distribution, and kME rankings were visualized, and gene-module assignments were exported for downstream interpretation and reproducibility.

### Cell–cell communication analysis

Cell–cell communication analysis was performed using the CellChat (90) package on the *Lama2/Esr1* neuronal population. The dataset was stratified by reproductive condition (Virgin and Lactating), and communication networks were inferred independently for each group using a curated mouse ligand–receptor database. Intercellular signaling was reconstructed by identifying overrepresented ligand–receptor interactions and estimating communication probabilities between neuronal subtypes. Interaction networks were aggregated at the pathway level to characterize global communication structure and signaling strength within each condition.

To enable direct comparison, condition-specific networks were integrated into a unified framework, allowing the assessment of differences in interaction number, strength, and pathway activity between the datasets.

Differential communication analysis was used to identify pathways exhibiting condition-dependent changes in signaling, as well as shifts in the relative contribution of specific neuronal populations. Functional organization of signaling networks was further explored by grouping pathways based on similarity (*k* = 4) in interaction patterns, enabling the identification of coordinated communication programs. Incoming and outgoing signaling patterns were analyzed to define the relative roles of neuronal subtypes as signal sources or targets, and to uncover state-dependent changes in network organization.

### NEUROeSTIMator activity scoring and cluster-level comparisons

Predicted neuronal activity scores were quantified at single-cell resolution using NEUROeSTIMator (40). Cells with missing predicted activity values were excluded prior to statistical testing. For the primary comparison, the *Gm33782/4933425B07Rik* cluster was tested against all remaining neuronal clusters combined using a two-sided Wilcoxon rank-sum test. In addition, cluster-wide activity profiles were summarized by computing the mean predicted activity per cluster and visualized as mean ± SEM to facilitate ranking across neuronal subtypes.

To test whether the subcluster differed from each neuronal cluster individually, pairwise two-sided Wilcoxon rank-sum tests were performed between the target cluster and each comparator cluster. Resulting *p* values were corrected for multiple comparisons using the Benjamini– Hochberg (BH) procedure, and adjusted significance levels were reported. Distributions were visualized using violin and box plots with jittered single-cell points, and pair-wise comparisons were displayed in faceted panels with BH-adjusted significance annotations.

### Trajectory inference and pseudotime analysis

Trajectory inference and pseudotime analysis were performed using Monocle3 (v1.3) to investigate transcriptional dynamics within MPOA neuronal populations (91). The annotated neuronal subset from the Seurat object was converted into a cell_data_set (CDS), preserving raw count matrices and cell-level metadata. Precomputed UMAP embeddings were retained as the low-dimensional representation for trajectory reconstruction, and cells were organized within this space to infer a principal trajectory structure.

Pseudotime ordering was established by selecting cells from the *Gm33782*/*4933425B07Rik* cluster enriched in the lactating condition as the root state, representing the initial transcriptional condition along the inferred trajectory. To identify genes exhibiting dynamic expression along pseudotime, a graph-based autocorrelation approach was applied to quantify the spatial dependency of gene expression patterns along the trajectory. Genes were ranked based on their degree of autocorrelation, and statistical significance was determined using multiple testing correction. Genes with adjusted *q*-values < 0.05 were considered significantly associated with pseudotime dynamics and were used for downstream analyses and visualization.

### SCENIC regulatory network inference

Gene regulatory network analysis was performed using the SCENIC framework to identify transcription factor–driven regulatory programs in MPOA neurons (92). Single-cell expression data processed in Seurat were exported to loom format and analyzed using pySCENIC (v0.12.1) executed within a Docker container (aertslab/pyscenic:0.12.1).

Candidate transcription factor–target gene interactions were first inferred using GRNBoost2, restricted to a curated list of mouse transcription factors. These co-expression modules were subsequently refined through cisTarget motif enrichment analysis using mouse motif ranking databases based on the mm10 genome assembly, thereby enabling the pruning of indirect interactions and the definition of high-confidence regulons. Regulon activity was then quantified in each cell using the AUCell algorithm, which computes the enrichment of regulon target genes among the highest expressed genes within individual transcriptomes.

### Enrichment analysis and IUPHAR classification

Functional enrichment analyses were performed to identify biological processes and signaling pathways associated with gene sets derived from differential expression and regulatory network analyses (93). Gene Ontology (GO), Kyoto Encyclopedia of Genes and Genomes (KEGG), and Reactome pathway enrichment analyses were conducted using the clusterProfiler (4.18.4) and ReactomePA (1.54.0) packages in R. Gene identifiers were mapped to Entrez IDs using the org.Mm.eg.db (3.22.0) annotation database. For Gene Ontology analysis, enrichment was evaluated across the Biological Process (BP), Molecular Function (MF), and Cellular Component (CC) categories. Background gene sets consisted of all genes detected in the single-cell dataset after quality control filtering. Enrichment significance was assessed using a hypergeometric test and resulting p-values were adjusted for multiple testing using the Benjamini–Hochberg false discovery rate (FDR) correction. Pathways with adjusted *q*-values < 0.05 were considered significantly enriched.

To further characterize the signaling landscape of *Pmch* neuronal populations, receptor genes identified in each subtype were annotated according to the International Union of Basic and Clinical Pharmacology (IUPHAR) classification system (94). Receptors were grouped into major families, including G protein–coupled receptors, ion channels, and enzyme-linked receptors, enabling systematic comparison of receptor repertoires between the *Tmem45a*/*Ror1* and *Gm33782*/*4933425B07Rik* populations. This integrative approach provides a pharmacologically informed framework to interpret potential ligand–receptor interactions and highlights subtype-specific signaling pathways associated with the molecular specialization of *Pmch* neurons within the MPOA.

## Supplementary Table Descriptions

**Table S1. Cluster-specific marker genes for major MPOA cell types identified using scType**. This tables lists the marker genes defining the principal cell populations identified in the MPOA using the scType annotation framework. For each cluster, genes are ranked according to log_2_ fold change, adjusted *p*-value, and the proportion of expressing cells. These markers support the annotation of major neuronal and non-neuronal cell types present in the dataset.

**Table S2. Marker genes defining neuronal subclusters within the *Lama2*/*Esr1* population**. This table lists genes enriched in transcriptionally distinct neuronal subclusters identified within the *Lama2*/*Esr1* neuronal population. For each gene, the table reports log_2_ fold change, adjusted *p*-value, and the proportion of expressing cells. These markers define the molecular identity of each subpopulation and reveal transcriptional heterogeneity within *Lama2*/*Esr1* neurons.

**Table S3. Marker genes defining the two *Pmch* neuronal populations within the MPOA *Lama2*/*Esr1* cluster**.

This table presents genes enriched in the two transcriptionally distinct *Pmch* neuronal populations identified in this study, defined by the molecular signatures *Tmem45a*/*Ror1* and *Gm33782*/*4933425B07Rik*. Shared and population-specific markers are listed to highlight both common molecular features and transcriptional differences between the two populations.

**Table S4. IUPHAR-based receptor annotation in *Pmch* neuronal subtypes**.

This table lists receptor genes expressed in the two *Pmch* neuronal populations defined by the molecular signatures *Tmem45a*/*Ror1* and *Gm33782*/*4933425B07Rik*. Receptors are classified according to the IUPHAR database and grouped by receptor family and subtype, providing a pharmacological overview of the signaling landscape associated with each population.

**Table S5. Intersection of marker genes between the two *Pmch* neuronal populations**. This table reports the overlap between marker genes identified in the *Tmem45a*/*Ror1* and *Gm33782*/*4933425B07Rik* neuronal populations, highlighting genes shared by both subtypes.

**Table S6. hdWGCNA modules identified in *Lama2*/*Esr1* neuronal subclusters**. This table lists gene co-expression modules identified by hdWGCNA within *Lama2*/*Esr1* neuronal subclusters. For each module, the table includes module assignment and the genes belonging to each module, defining coordinated transcriptional programs associated with specific neuronal states.

**Table S7. Intersection between hdWGCNA modules and marker genes of the two *Pmch* neuronal populations**.

This table shows the overlap between genes assigned to hdWGCNA modules and marker genes defining the two *Pmch* neuronal populations (*Tmem45a*/*Ror1* and *Gm33782*/*4933425B07Rik*), identifying module-associated genes potentially contributing to the molecular identity of these populations.

**Table S8. Functional enrichment of genes shared between hdWGCNA modules and *Pmch* markers**. This table summarizes Gene Ontology (GO), KEGG, and Reactome enrichment analyses performed on genes shared between hdWGCNA modules and markers of the two *Pmch* neuronal populations.

**Table S9. NeuroeSTIMator activity scores across *Lama2*/*Esr1* neuronal subpopulations**. This table reports NeuroeSTIMator-derived neuronal activity scores across all *Lama2*/*Esr1* neuronal subclusters, including the two *Pmch* populations, allowing comparison of predicted neuronal activity states across subpopulations.

**Table S10. CellChat-inferred intercellular communication network**.

This table summarizes ligand–receptor interactions inferred by CellChat across MPOA neuronal populations, including interaction strength, signaling pathways, and source–target relationships for all cell types.

**Table S11. Transcription factor regulons identified by SCENIC analysis**.

This table lists transcription factor (TF) regulons inferred using SCENIC, including TF identity, predicted target genes, and regulon activity scores across MPOA neuronal populations.

**Table S12. Pseudotime trajectory metrics across *Lama2/Esr1* neuronal subclusters**. This table provides pseudotime values assigned to individual *Lama2*^*+*^*/Esr1*^*+*^ neurons along the inferred trajectory, including cells belonging to the two *Pmch* neuronal populations.

**Table S13. Intersection between predicted *Pmch* regulators and *Pmch* neuronal population markers**. This table reports the overlap between TFs identified through regulatory network inference and genes predicted to regulate *Pmch*, alongside marker genes defining the two *Pmch* neuronal populations.

**Table S14. Functional enrichment of transcription factors predicted to regulate *Pmch***. This table summarizes Gene Ontology (GO), KEGG, and Reactome enrichment analyses performed on TFs predicted to regulate *Pmch*, derived from the intersection between TF regulons and marker genes of the two *Pmch* neuronal populations (*Tmem45a*/*Ror1* and *Gm33782/4933425B07Rik*).

**Table S15. Gene panel used for single-cell RT–qPCR (*Biomark HD* system)**. This table lists the genes included in the 48-gene panel used for single-cell expression profiling using the *Biomark HD* microfluidic platform. The panel was designed to capture neurotransmitter identity, neuropeptidergic signaling, and hormone responsiveness in MPOA neurons. For each gene, the table includes the assay ID and functional category.

**Fig. S1.**
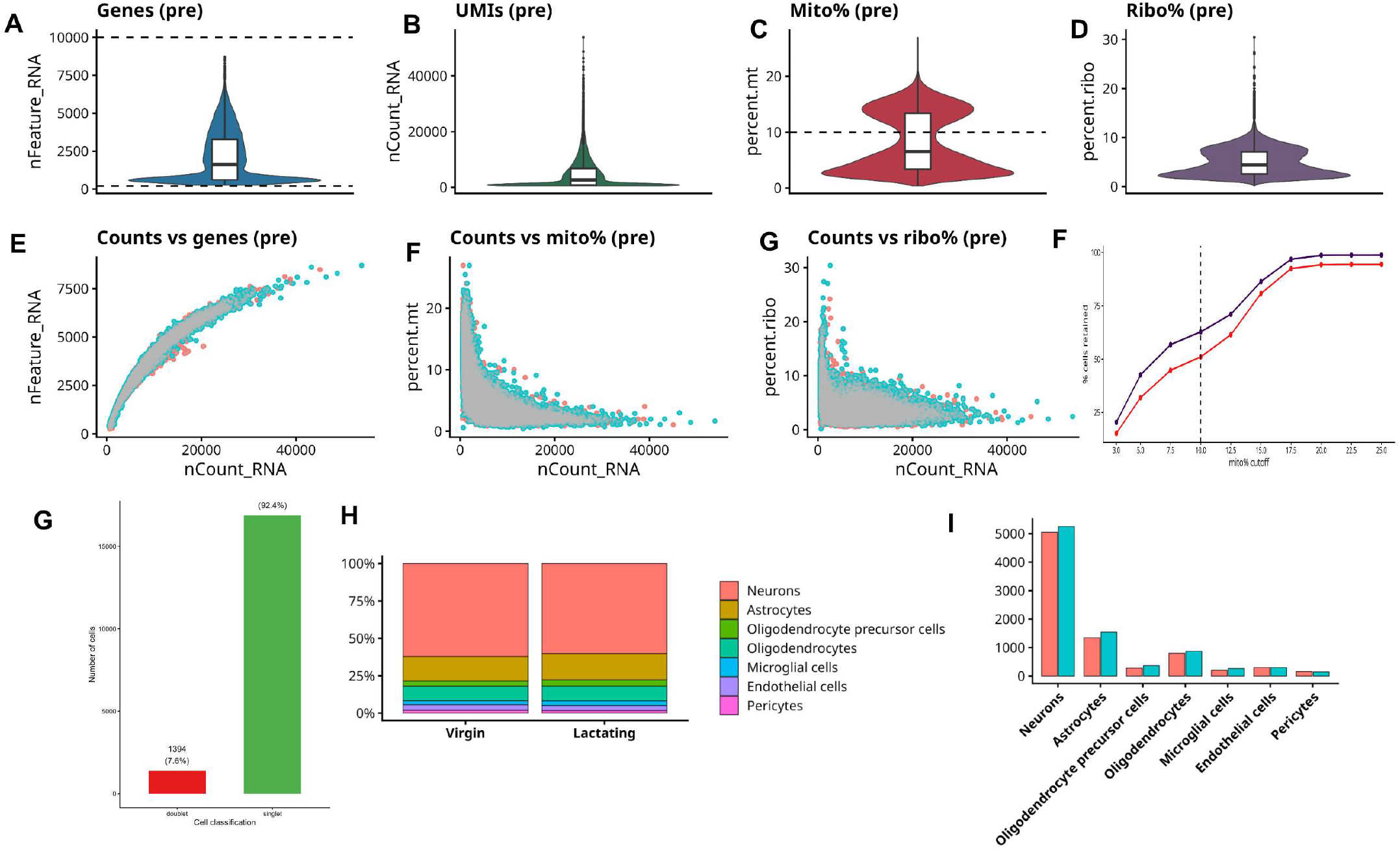
Quality control metrics for single-cell RNA-seq data from the MPOA dataset. **A–D**. Violin plots showing the distribution of detected genes (*nFeature_RNA*), total UMI counts (*nCount_RNA*), mitochondrial transcript percentage (mito%), and ribosomal transcript percentage (ribo%) prior to filtering. Dashed lines indicate the thresholds used for quality control. **E–G**. Scatter plots illustrating the relationships between UMI counts and detected genes, mitochondrial content, and ribosomal content across cells before filtering. **F**. Saturation curve showing the proportion of retained cells across increasing mitochondrial thresholds used to define the filtering cutoff. **G**. Bar plot showing the classification of droplets into singlets and doublets following doublet detection analysis. **H**. Relative proportion of major cell types identified in the dataset across virgin and lactating conditions. I. Absolute number of cells per major cell type across conditions, including neurons, astrocytes, oligodendrocyte precursor cells, oligodendrocytes, microglial cells, endothelial cells, and pericytes

**Fig. S2.**
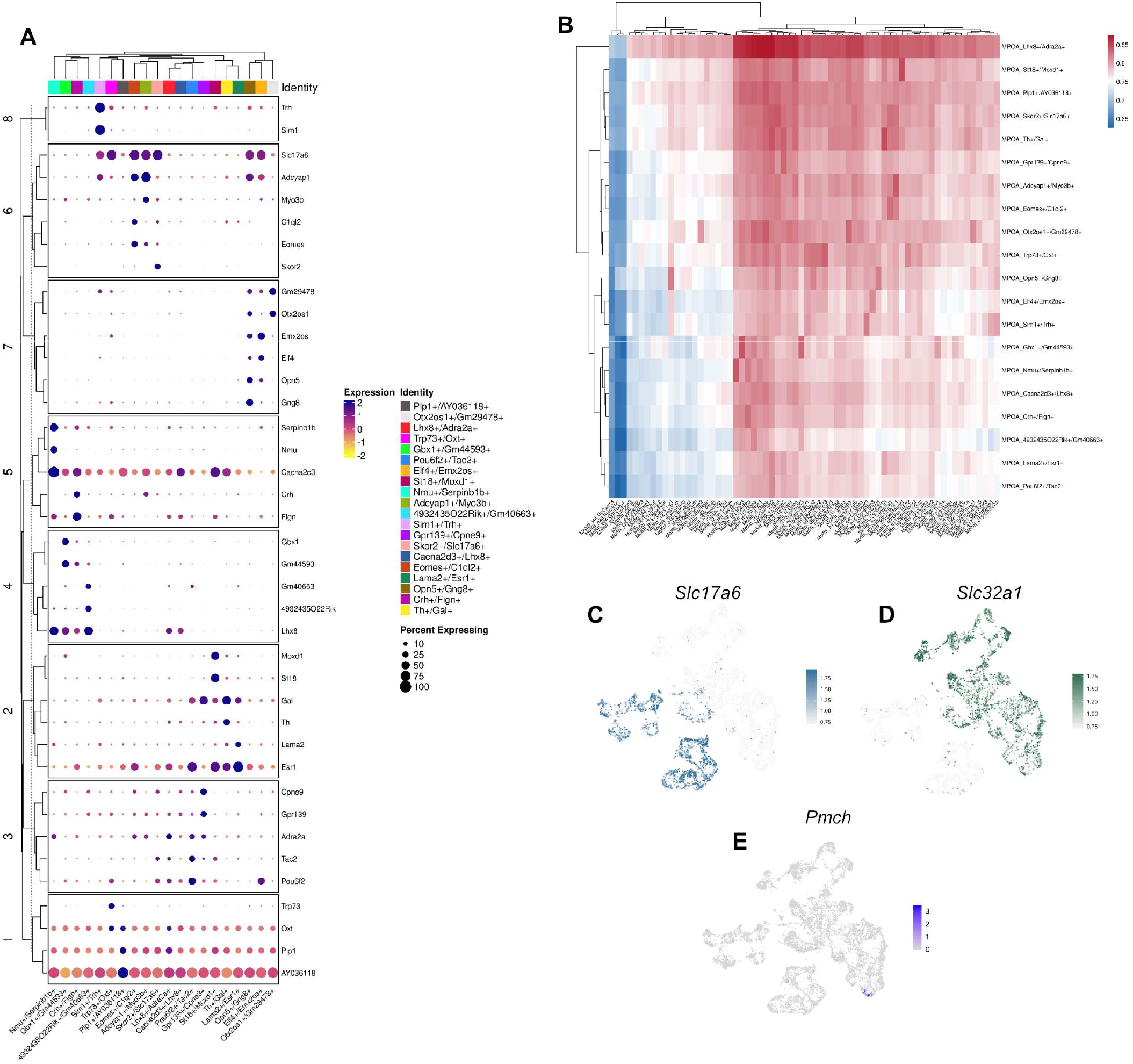
Molecular identity and neurotransmitter profile of MPOA neuronal subclusters. **A**. Dot plot showing hierarchical clustering of marker gene expression across neuronal subclusters in the MPOA. Distinct transcriptional signatures define each cluster, including enrichment of neuropeptides (*Gal, Nmu, Crh, Trh*), transcription factors (*Sim1, Eomes*), and region-specific markers, highlighting the molecular heterogeneity of MPOA neurons. **B–C**. Feature plots showing expression of neurotransmitter-related genes across neuronal populations. Expression of *Slc17a6* identifies glutamatergic neurons, whereas *Slc32a1* marks GABAergic populations, indicating that the majority of MPOA neurons exhibit an inhibitory phenotype. **D**. Feature plot of *Pmch* expression reveals a spatially restricted distribution within a subset of neuronal clusters, consistent with selective enrichment of MCH neurons within specific MPOA subpopulations.

**Fig. S3.**
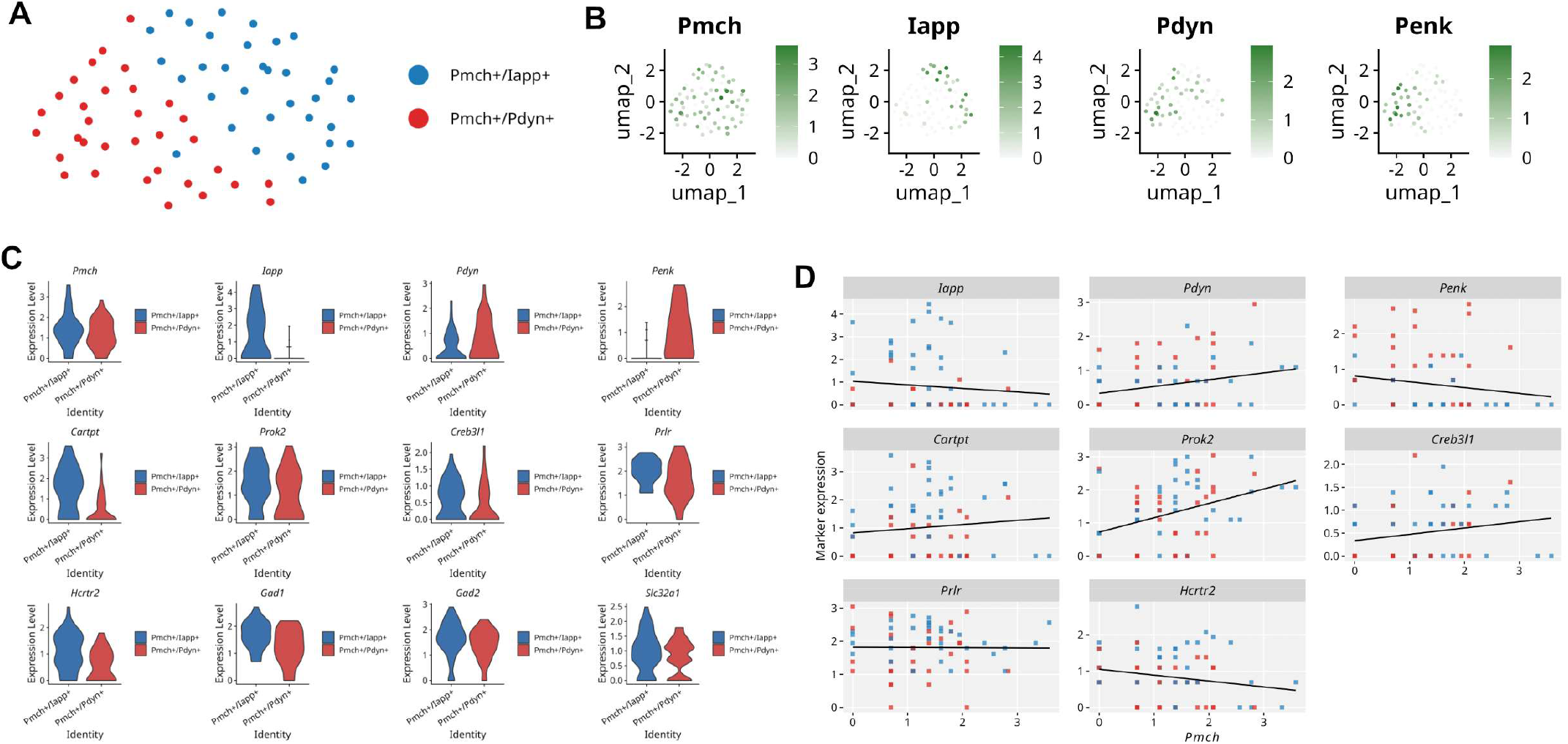
Subclustering and transcriptional characterization of *Pmch* neuronal populations following focused reanalysis of *Pmch*-expressing cells (logCPM > 1). **A**. UMAP representation of the subsetted *Pmch*-expressing neuronal population, revealing two transcriptionally distinct subclusters characterized by preferential enrichment of *Iapp* (*Pmch/Iapp*) or *Pdyn* (*Pmch/Pdyn*) expression. **B**. Feature plots showing the distribution and expression intensity of *Pmch, Iapp, Pdyn*, and *Penk* across the re-clustered *Pmch* neuronal populations. **C**. Violin plots comparing expression levels of representative marker genes between *Pmch/Iapp* and *Pmch/Pdyn* neuronal subpopulations, including *Pmch, Iapp, Pdyn, Penk, Cartpt, Prok2, Creb3l1, Prlr, Hcrtr2, Gad1, Gad2*, and *Slc32a1*. **D**. Scatterplot analyses showing the relationship between *Pmch* expression and representative marker genes across individual cells within the subsetted *Pmch* neuronal populations. Linear regression trends indicate distinct transcriptional associations between *Pmch* and neuropeptidergic markers, supporting the existence of transcriptionally heterogeneous *Pmch* neuronal subtypes within the MPOA.

**Fig. S4.**
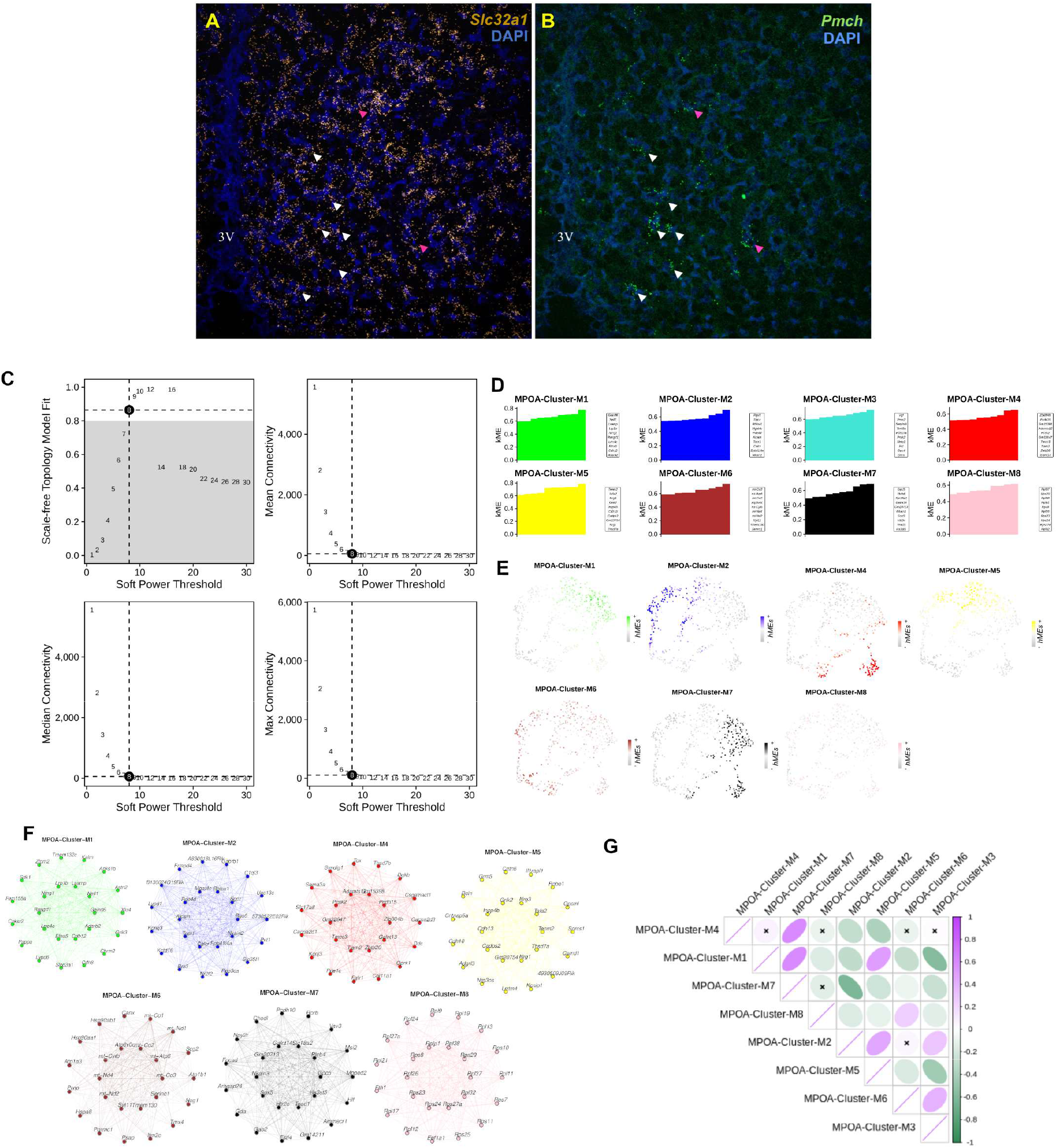
Validation of GABAergic identity and construction of gene co-expression networks in MPOA neuronal populations. **A–B**. Fluorescence *in situ* hybridization (RNAscope) showing expression of *Slc32a1* (A, yellow) and *Pmch* (B, green) in the MPOA, with DAPI nuclear staining (blue). White arrowheads indicate neurons co-expressing *Slc32a1* and *Pmch*, confirming the GABAergic identity of *Pmch*-expressing neurons, while pink arrowheads indicate single-positive cells. The third ventricle (3V) is indicated for anatomical reference. **C**. Soft-thresholding power analysis used to determine the optimal parameter for weighted gene co-expression network construction. Plots show the scale-free topology model fit, mean connectivity, median connectivity, and maximum connectivity across candidate soft power thresholds. The selected soft power is indicated by dashed lines. **D**. Gene membership (kME) distributions for hdWGCNA modules (M1– M8). Genes are ranked according to module membership (kME), highlighting those most strongly associated with each module eigengene. **E**. UMAP projections showing module eigengene expression across single cells for each hdWGCNA module (M1–M8), illustrating the spatial distribution of module activity across transcriptionally defined neuronal populations in the MPOA. **F**. Gene co-expression network representations for each hdWGCNA module (M1–M8). Nodes represent genes and edges represent co-expression relationships, highlighting the internal connectivity structure of each module. **G**. Module–cluster correlation matrix showing the association between hdWGCNA modules and MPOA neuronal clusters. Color intensity and ellipse orientation indicate the strength and direction of the correlation, while crosses denote non-significant associations

**Fig. S5.**
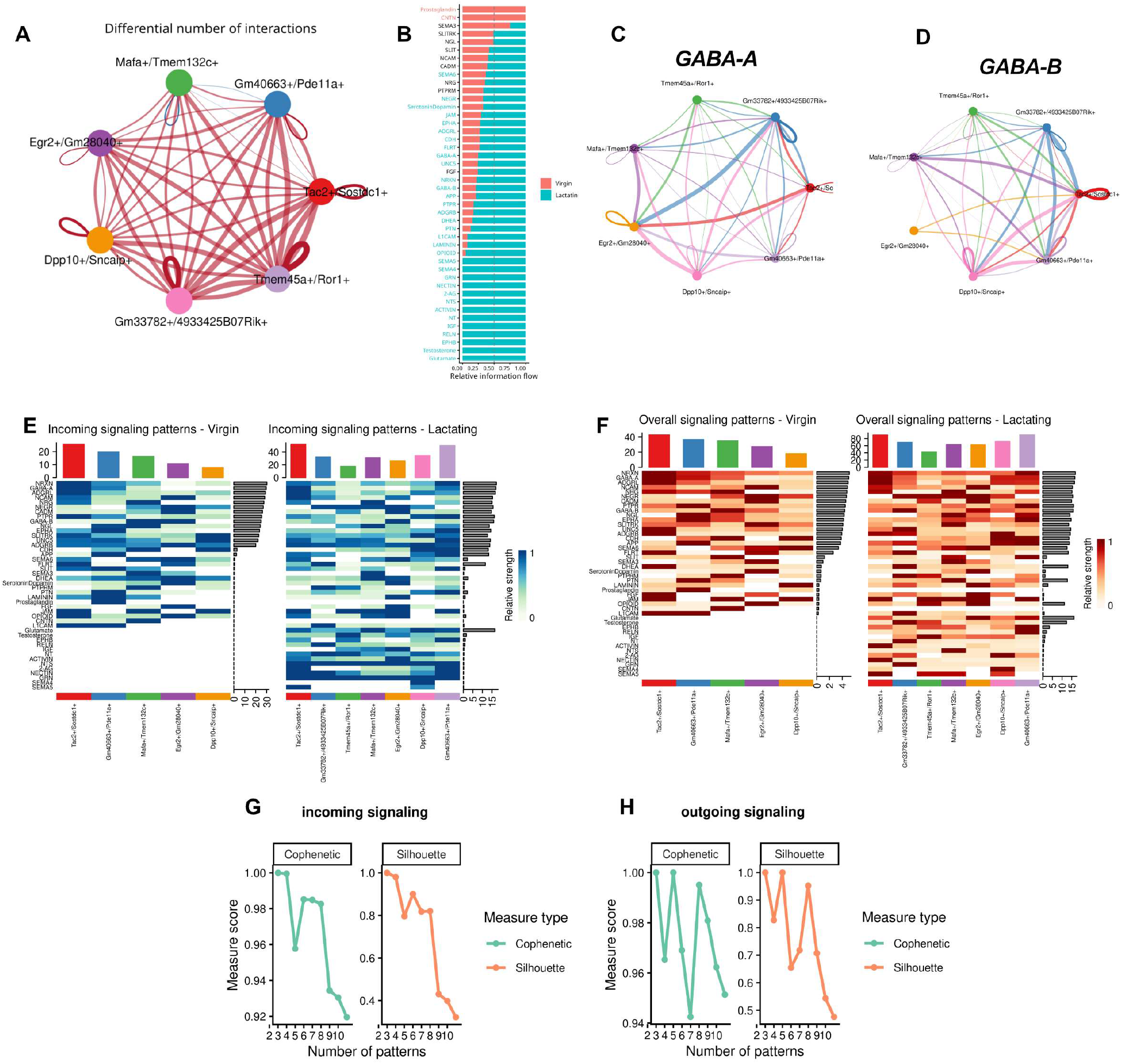
State-dependent organization of MPOA^*Lama2/Esr1*^ communication networks. **A**. Differential number of inferred interactions across MPOA^*Lama2/Esr1*^ neuronal subtypes, highlighting changes in network connectivity. **B**. Relative contribution of each neuronal population to overall information flow, comparing virgin and lactating conditions. **C–D**. Representative inhibitory signaling networks (GABA-A and GABA-B), illustrating interaction patterns among neuronal subtypes. **E**. Incoming signaling patterns in virgin and lactating states, showing condition-dependent differences in pathway distribution across neuronal populations. **F**. Global signaling patterns in virgin and lactating conditions, indicating increased pathway activity and coordination during lactation. (G–H) Pattern selection analysis for incoming **G**. and outgoing **H**. signaling, with Cophenetic and Silhouette metrics supporting the robustness of inferred communication programs.

**Fig. S6.**
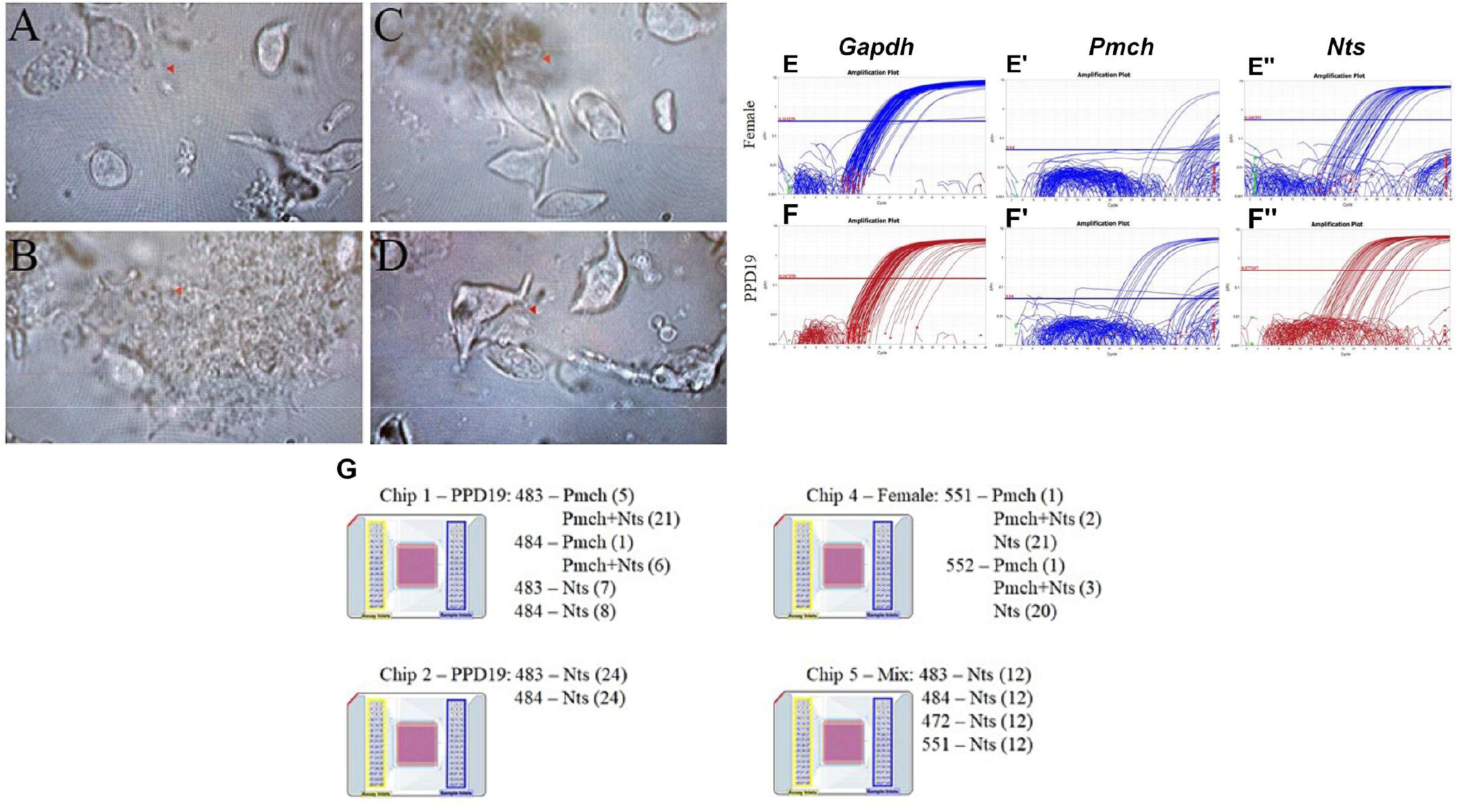
Single-cell isolation, RT-qPCR screening, and microfluidic gene expression profiling of MPOA GAD67-GFP neurons. **A–D**. Representative images of dissociated MPOA cells prior to sorting, illustrating cellular integrity and morphology following micropunch isolation and trituration. Arrowheads indicate individual viable neurons. **E–E″**. Representative RT-qPCR amplification curves for *Gapdh, Pmch*, and *Nts* in virgin female animals, showing robust detection of *Gapdh* and *Nts*, with absence of *Pmch* expression. **F–F″**. Representative RT-qPCR amplification curves for *Gapdh, Pmch*, and *Nts* in lactating animals (PPD19), demonstrating selective expression of *Pmch* in the lactating condition. **G**. Schematic representation of microfluidic chip loading strategy (48.48 dynamic arrays, *Biomark HD* system), indicating distribution of single cells across experimental groups and neurochemical categories (*Pmch, Nts*, and *Pmch/Nts*).

**Fig. S7.**
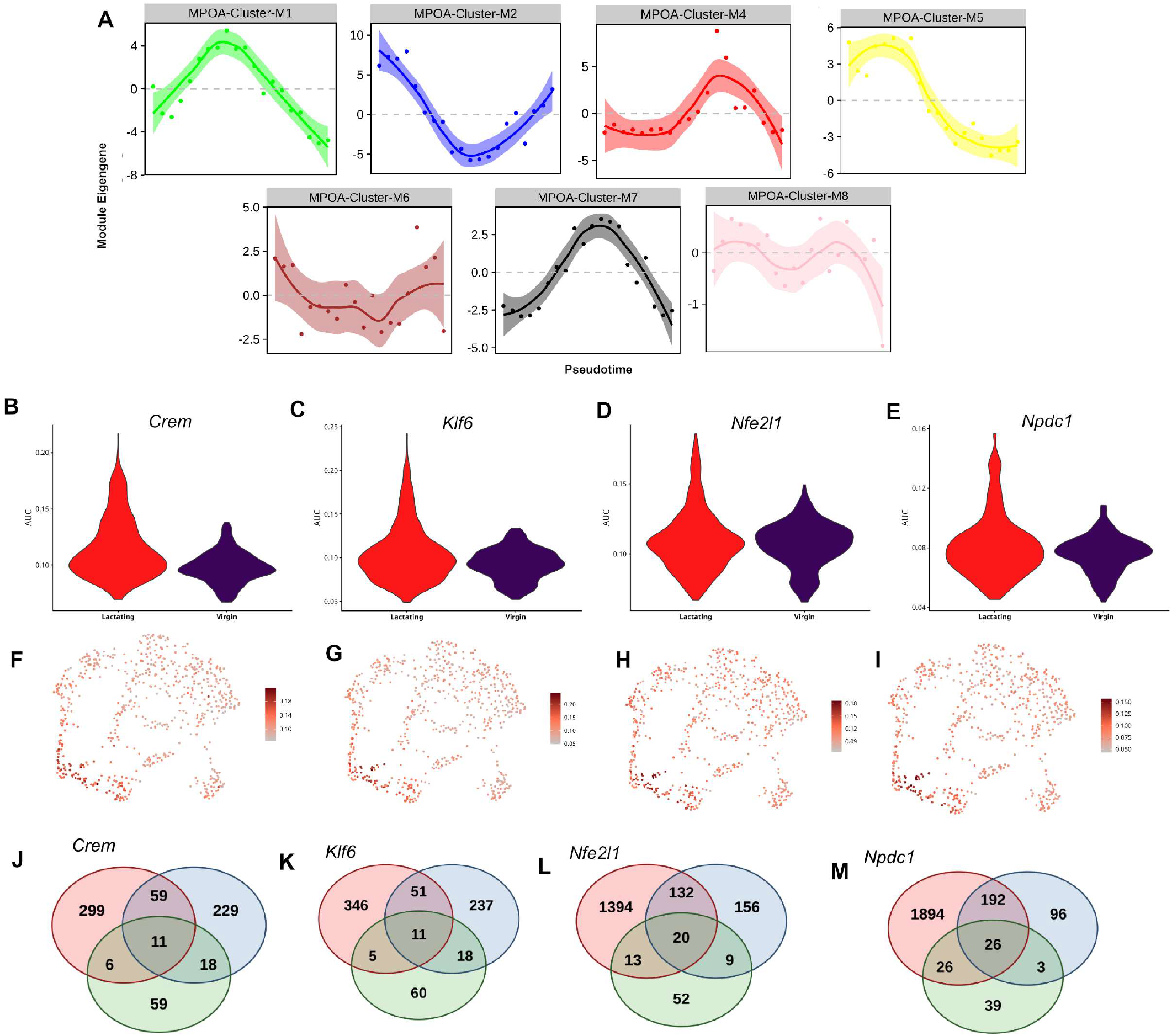
Transcription factor activity and target gene intersections associated with *Pmch* neuronal populations. **A**. Module eigengene dynamics across pseudotime for multiple MPOA-associated co-expression modules (M1–M8), revealing distinct temporal activation patterns and indicating that transcriptional programs are differentially engaged along the trajectory. **B–E**. Violin plots showing regulon activity scores (AUC) for additional transcription factors (*Crem, Klf6, Nfe2l1, Npdc1*) across physiological conditions, indicating increased regulatory activity in lactating animals compared with virgins. **F–I**. UMAP visualization of transcription factor activity scores demonstrates spatial distribution of regulon activity across the neuronal manifold, supporting region-specific regulatory engagement. **J–M**. Intersection analysis of transcription factor target genes shows overlap between regulon targets, and cluster-specific markers, identifying shared and unique downstream regulatory programs for each transcription factor.

